# Evolution of scapula size and shape in Carnivora: locomotor habits and differential shape scaling

**DOI:** 10.1101/2022.08.26.505396

**Authors:** Eloy Gálvez-López, Adrià Casinos

## Abstract

The effect of size, phylogeny, and locomotor habit, on shape was tested in 213 scapulas from 101 carnivoran species using 3D geometric morphometric methods. The sampled species spanned the whole size range and locomotor patterns in Carnivora. The results of the present study indicate that, in this order, scapula shape responds to the complex interaction of allometric, phylogenetic, and functional effects. Furthermore, evidence for differential scaling in the shape of the carnivoran scapula was found, which might be related to scaling differences among carnivoran families. Additionally, most allometric shape variation in the carnivoran scapula was related to size changes along phyletic lines. Locomotor-related shape differences were assessed using canonical variate analysis. Most locomotor habits could be significantly separated from each other based on scapula shape, although high misclassification rates were obtained when comparing semiarboreal and semifossorial carnivorans to other locomotor types. Locomotor indicators in the scapula shape of extant carnivorans seemed independent of size or shared ancestry and could be related to muscular function. These locomotor indicators were then used to infer the locomotor habits of several internal nodes of the carnivoran phylogeny, whose scapular size and shape was reconstructed using weighted square-change parsimony. According to scapula size and shape, the carnivoran ancestor was a medium-sized scansorial animal (i.e., it spent most of its time on the ground, but was a good climber).

## Introduction

The therian scapula consists of a slightly concavoconvex bone plate (scapular blade) whose lateral surface is divided by the scapular spine in two fossae (supraspinous and infraspinous). The medial surface of the scapular blade is known as subscapular fossa. The scapular spine is a roughly flat plate of bone extending almost perpendicularly to the scapular blade. The spine broadens distally into the acromion, which can present a ventral projection (hamatus process) and a caudal extension (suprahamatus process or metacromion). The scapula articulates distally with the proximal humerus at the glenoid cavity, which is separated from the rest of the scapular blade by a strangulation known as scapular neck. Finally, in some species a coracoid process may arise cranio-medially to the glenoid. As stated by Monteiro & Abe (1999), the therian scapula is a complex morphological structure (Atchley & Hall, 1991), since it derives from two ossification centers with different ontogenetic origin: the scapular plate and the coracoid plate (Goodrich, 1930).

Scapular morphology arises from the combination of phylogenetic history and functional requirements. The scapula is both an element of the thoracic girdle, providing insertion to the muscles connecting the forelimb to the rest of the body (Lessertisseur & Saban, 1967), and a functional element of the forelimb, being the most propulsive segment during locomotion (Fischer et al., 2002). Thus, its functional requirements include both shoulder stabilization and scapular mobility. The relative importance of those functions, however, surely varies among specialized locomotor patterns. For instance, a slow arboreal species would require stronger stabilization to avoid shoulder dislocation while navigating the three-dimensional pathways of the canopy, whereas larger scapular mobility in the parasagittal plane would increase the performance of a fast cursorial species (sensu Stein & Casinos, 1997) by allowing larger strides. Furthermore, differences in body size are expected to also influence scapular morphology, since body size plays a major role determining the biomechanics of locomotion (Schmidt-Nielsen, 1984; Alexander, 2002; Biewener, 2003). Thus, the scapula is considered particularly suited for studies on ecomorphology and morphological evolution (Astúa, 2009).

Most of the previous ecomorphological studies on the therian scapula focus on functional anatomy, relating sets of anatomical characters to particular locomotor habits in the studied groups (Davis, 1949; Maynard Smith & Savage, 1956; Lehmann, 1963; Ashton et al., 1965; Müller, 1967; Oxnard, 68; Roberts, 1974;Taylor, 1974; English, 1977; Taylor, 1997; Argot, 2001; Seckel & Janis, 2008). The development of geometric morphometrics (GM) methods in the 1990s allowed the separate analysis of shape and size in morphological studies. To date, scapular shape has been studied mainly in rodents (Swiderski, 1993; Morgan, 2009), dolphins (Smith et al., 1994), xenarthrans (Monteiro & Abe, 1999; Monteiro, 2000), primates (Young, 2004; Taylor & Slice, 2005; Young, 2008), and didelphids (Astúa, 2009). According to these studies, scapular shape is heavily influenced by phylogeny, while the amount of scapular shape variation that can be attributed to locomotor differences varies among groups. Regarding size, a significant allometric effect on scapular shape was found in didelphids (Astúa, 2009), but not in xenarthrans (Monteiro & Abe, 1999) or caviomorph rodents (Morgan, 2009).

The order Carnivora currently comprises over 280 species in 16 families, and constitutes a well-defined monophyletic group (Wilson & Mittermeier, 2009; Nyakatura & Bininda-Emonds, 2012). Carnivorans present one of the widest locomotor diversities among mammals, which makes them perfect subjects for ecomorphological studies (e.g. Oxnard, 1968; Taylor, 1974; Van Valkenburgh, 1985, 1987; Iwaniuk et al., 1999). Furthermore, they span a size range of four orders of magnitude (from less than 0.1 kg in the least weasel (*Mustela nivalis*) to well over two tonnes in elephant seals (*Mirounga sp*.)), which provides a solid base from which to test for allometric effects. Several researchers have analyzed bone shape in Carnivora using GM methods, particularly on the cranium and mandible (e.g. Goswami, 2006; Meloro et al., 2008; Figueirido et al., 2010) and, to a lesser extent, the long bones of the limbs (Schutz & Guralnick, 2007; Walmsley et al., 2012; Fabre et al., 2013*a*,*b*; Martín-Serra et al., 2014). However, despite the remarkable shape variability of the scapula and its substantial biomechanical importance, only the work of Martín-Serra et al. (2014) included this bone in their samples, and then only to the distalmost scapula (acromion and coracoglenoid region) plus teres major process.

Thus, the first aim of the present study is to explore the shape variability of the carnivoran scapula, and to quantify to what extent, if any, phylogenetic history and size differences affect scapula shape. The second aim is to assess whether indicators of particular locomotor habits (e.g. climbing, swimming) can be identified in the carnivoran scapula, and to characterize them in case they do. Finally, should these locomotor indicators exist, the locomotor type of the carnivoran ancestor could be inferred according to the reconstructed evolution of scapula shape in this clade.

## Material and Methods

The sample consisted of 213 scapulas from 101 species of Carnivora (Table 1), representing all extant families but Odobenidae and Prionodontidae (Wozencraft, 2005; Wilson & Mittermeier, 2009). Furthermore, the sampled species span the whole size range of the order, as well as all locomotor habits in Carnivora. Following our previous research (Gálvez-López & Casinos, 2022*a*, *b*), the locomotor specialization of each species (i.e., its main locomotor habit according to the literature) was summarized in a series of locomotor categories (Table 2). The phylogeny proposed by Nyakatura & Bininda-Emonds (2012) was used, although with slight modifications (see Supplementary Information of Gálvez-López (2021) for a detailed description). A pruned version including the species analyzed in this study can be found in Figure 1.

**Table 1.** Measured species. For each species, the table shows the abbreviation used in the text and figures (abbr.), the number of digitized specimens (n), and the assigned locomotor type category (loctyp). See Table 2 for a description of locomotor type categories and their abbreviations.

| species | abbr. | n | loctyp | species | abbr. | n | loctyp | species | abbr. | n | loctyp |
| --- | --- | --- | --- | --- | --- | --- | --- | --- | --- | --- | --- |
| Canidae |  |  |  |  |  |  |  |  |  |  |  |
| *Canis aureus* | Cau | 1 | terr | *Lycalopex culpaeus* | Lcu | 1 | terr | *Vulpes chama* | Vch | 1 | terr |
| *Canis lupus* | Clu | 4 | terr | *Lycalopex gymnocercus* | Lgy | 2 | terr | *Vulpes vulpes* | Vvu | 11 | terr |
| *Chrysocyon brachyurus* | Cbr | 2 | terr | *Lycaon pictus* | Lpi | 2 | terr | *Vulpes zerda* | Vze | 1 | terr |
| Mustelidae |  |  |  |  |  |  |  |  |  |  |  |
| *Eira barbara* | Eba | 1 | sarb | *Lutra lutra* | Llu | 3 | saq | *Mustela erminea* | Mer | 2 | terr |
| *Enhydra lutris* | Elu | 1 | aq | *Lyncodon patagonicus* | Lpt | 2 | terr | *Mustela eversmannii* | Mev | 1 | terr |
| *Galictis cuja* | Gcu | 2 | terr | *Martes americana* | Mam | 1 | sarb | *Mustela lutreola* | Mlu | 1 | saq |
| *Galictis vittata* | Gvi | 1 | terr | *Martes foina* | Mfo | 18 | scan | *Mustela nivalis* | Mvi | 2 | terr |
| *Gulo gulo* | Ggu | 1 | scan | *Martes martes* | Mma | 3 | sarb | *Mustela putorius* | Mpu | 2 | terr |
| *Ictonyx striatus* | Ist | 1 | saq | *Martes zibellina* | Mzi | 1 | scan | *Neovison vison* | Nvi | 2 | saq |
| *Lontra felina* | Lfe | 2 | saq | *Meles meles* | Mme | 3 | sfos | *Pteronura brasiliensis* | Pbr | 1 | saq |
| *Lontra longicaudis* | Llo | 1 | saq | *Mellivora capensis* | Mca | 1 | sfos |  |  |  |  |
| *Lontra provocax* | Lpr | 2 | saq | *Melogale moschata* | Mmo | 1 | terr |  |  |  |  |
| Mephitidae |  |  |  |  |  |  |  |  |  |  |  |
| *Conepatus chinga* | Cch | 2 | sfos | *Spilogale gracilis* | Sgr | 2 | terr |  |  |  |  |
| Phocidae |  |  |  |  |  |  |  |  |  |  |  |
| *Hydrurga leptonyx* | Hle | 1 | aq | *Mirounga leonina* | Mle | 1 | aq | *Phoca vitulina* | Pvi | 1 | aq |
| Otariidae |  |  |  |  |  |  |  |  |  |  |  |
| *Arctocephalus australis* | Aau | 1 | aq | *Otaria flavescens* | Ofl | 2 | aq |  |  |  |  |
| *Arctocephalus gazella* | Aga | 1 | aq | *Zalophus californianus* | Zca | 2 | aq |  |  |  |  |
| Ailuridae |  |  |  |  |  |  |  | Nandiniidae |  |  |  |
| *Ailurus fulgens* | Afu | 7 | scan |  |  |  |  | *Nandinia binotata* | Nbi | 4 | sarb |
| Procyonidae |  |  |  |  |  |  |  |  |  |  |  |
| *Bassaricyon gabbii* | Bga | 1 | arb | *Nasua nasua* | Nna | 2 | scan | *Procyon lotor* | Plo | 3 | scan |
| *Bassariscus astutus* | Bas | 1 | scan | *Potos flavus* | Pfl | 2 | arb |  |  |  |  |
| *Nasua narica* | Nnr | 4 | scan | *Procyon cancrivorus* | Pca | 2 | scan |  |  |  |  |
| Ursidae |  |  |  |  |  |  |  |  |  |  |  |
| *Ailuropoda melanoleuca* | Ame | 1 | scan | *Ursus americanus* | Uam | 1 | scan | *Ursus maritimus* | Uma | 3 | terr |
| *Tremarctos ornatus* | Tor | 1 | scan | *Ursus arctos* | Uar | 4 | scan |  |  |  |  |
| Viverridae |  |  |  |  |  |  |  |  |  |  |  |
| *Arctictis binturong* | Abi | 3 | arb | *Genetta maculate* | Gma | 3 | sarb | *Viverra zibetha* | Vzi | 2 | terr |
| *Arctogalidia trivirgata* | Atr | 2 | arb | *Paradoxurus hermaphroditus* | Phe | 2 | arb | *Viverricula indica* | Vin | 3 | scan |
| *Civettictis civetta* | Cci | 2 | terr | *Poiana richardsoni* | Pri | 1 | sarb |  |  |  |  |
| *Genetta genetta* | Gge | 6 | scan | *Viverra tangalunga* | Vta | 3 | terr |  |  |  |  |
| Herpestidae |  |  |  |  |  |  |  |  |  |  |  |
| *Atilax paludinosus* | Apa | 1 | saq | *Herpestes ichneumon* | Hic | 3 | terr | *Urva edwardsii* | Ued | 2 | terr |
| *Crossarchus obscurus* | Cob | 1 | terr | *Ichneumia albicauda* | Ial | 1 | terr | *Urva javanica* | Uja | 1 | terr |
| *Galerella sanguinea* | Gsa | 1 | terr | *Suricata suricatta* | Ssu | 2 | sfos |  |  |  |  |
| Eupleridae |  |  |  |  |  |  |  |  |  |  |  |
| *Cryptoprocta ferox* | Cfe | 2 | sarb | *Galidia elegans* | Gel | 2 | scan | *Salanoia concolor* | Sco | 2 | scan |
| *Fossa fossana* | Ffo | 1 | terr | *Mungotictis decemlineata* | Mde | 1 | scan |  |  |  |  |
| Hyaenidae |  |  |  |  |  |  |  |  |  |  |  |
| *Crocuta Crocuta* | Ccr | 1 | terr | *Hyaena hyaena* | Hhy | 1 | terr | *Proteles cristatus* | Pcr | 1 | terr |
| Felidae |  |  |  |  |  |  |  |  |  |  |  |
| *Acinonyx jubatus* | Aju | 2 | scan | *Leopardus pardalis* | Lpa | 1 | scan | *Panthera leo* | Ple | 2 | scan |
| *Caracal aurata* | Cau | 1 | scan | *Leopardus tigrinus* | Lti | 1 | scan | *Panthera onca* | Pon | 2 | scan |
| *Felis silvestris* | Fsi | 2 | scan | *Leptailurus serval* | Lse | 2 | scan | *Panthera pardus* | Ppa | 3 | scan |
| *Herpailurus yaguaroundi* | Hya | 1 | scan | *Lynx lynx* | Lly | 1 | scan | *Panthera tigris* | Pti | 3 | scan |
| *Leopardus colocolo* | Lco | 1 | scan | *Lynx pardinus* | Lpd | 4 | scan | *Panthera uncia* | Pun | 2 | scan |
| *Leopardus geoffroyi* | Lge | 2 | scan | *Lynx rufus* | Lru | 1 | scan | *Puma concolor* | Pco | 2 | scan |

**Table 2.**
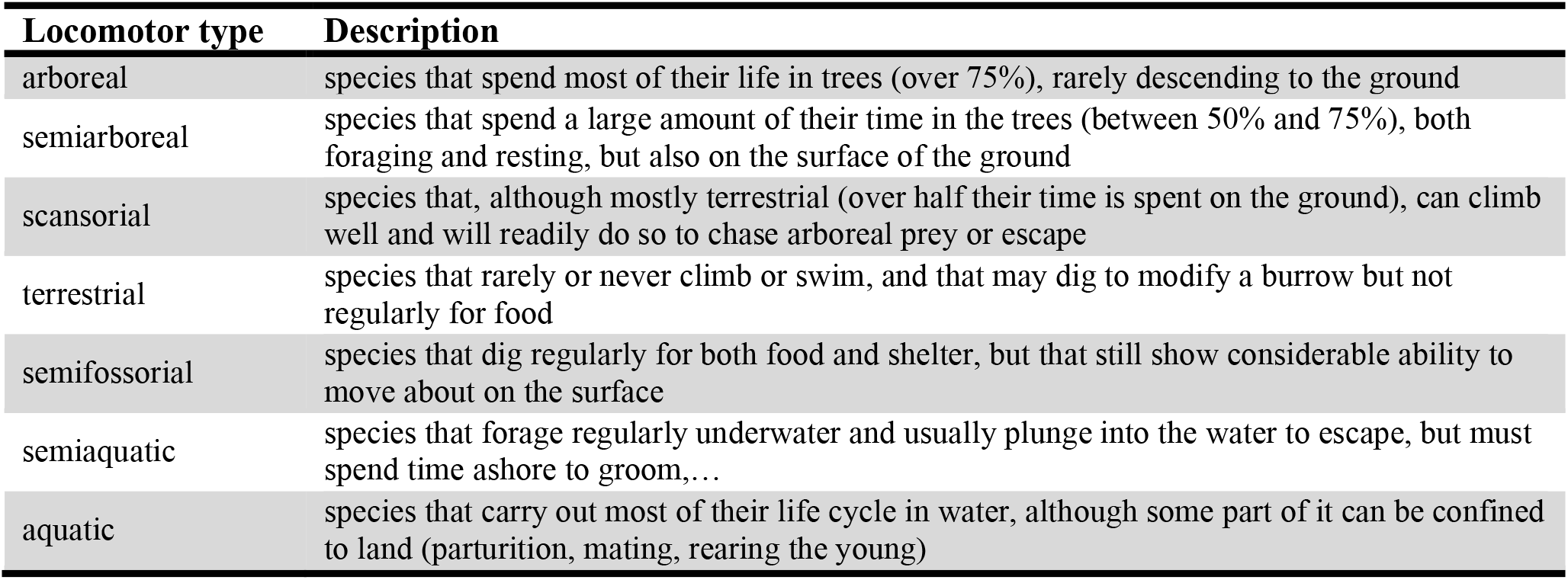
Locomotor type categories. The categories were adapted from previous works on the relationship between locomotor habit and forelimb morphology (Eisenberg, 1981; Van Valkenburgh, 1985, 1987).

| Locomotor type | Description |
| --- | --- |
| arboreal | species that spend most of their life in trees (over 75%), rarely descending to the ground |
| semiarboreal | species that spend a large amount of their time in the trees (between 50% and 75%), both foraging and resting, but also on the surface of the ground |
| scansorial | species that, although mostly terrestrial (over half their time is spent on the ground), can climb well and will readily do so to chase arboreal prey or escape |
| terrestrial | species that rarely or never climb or swim, and that may dig to modify a burrow but not regularly for food |
| semifossorial | species that dig regularly for both food and shelter, but that still show considerable ability to move about on the surface |
| semiaquatic | species that forage regularly underwater and usually plunge into the water to escape, but must spend time ashore to groom,... |
| aquatic | species that carry out most of their life cycle in water, although some part of it can be confined to land (parturition, mating, rearing the young) |

**Figure 1.**
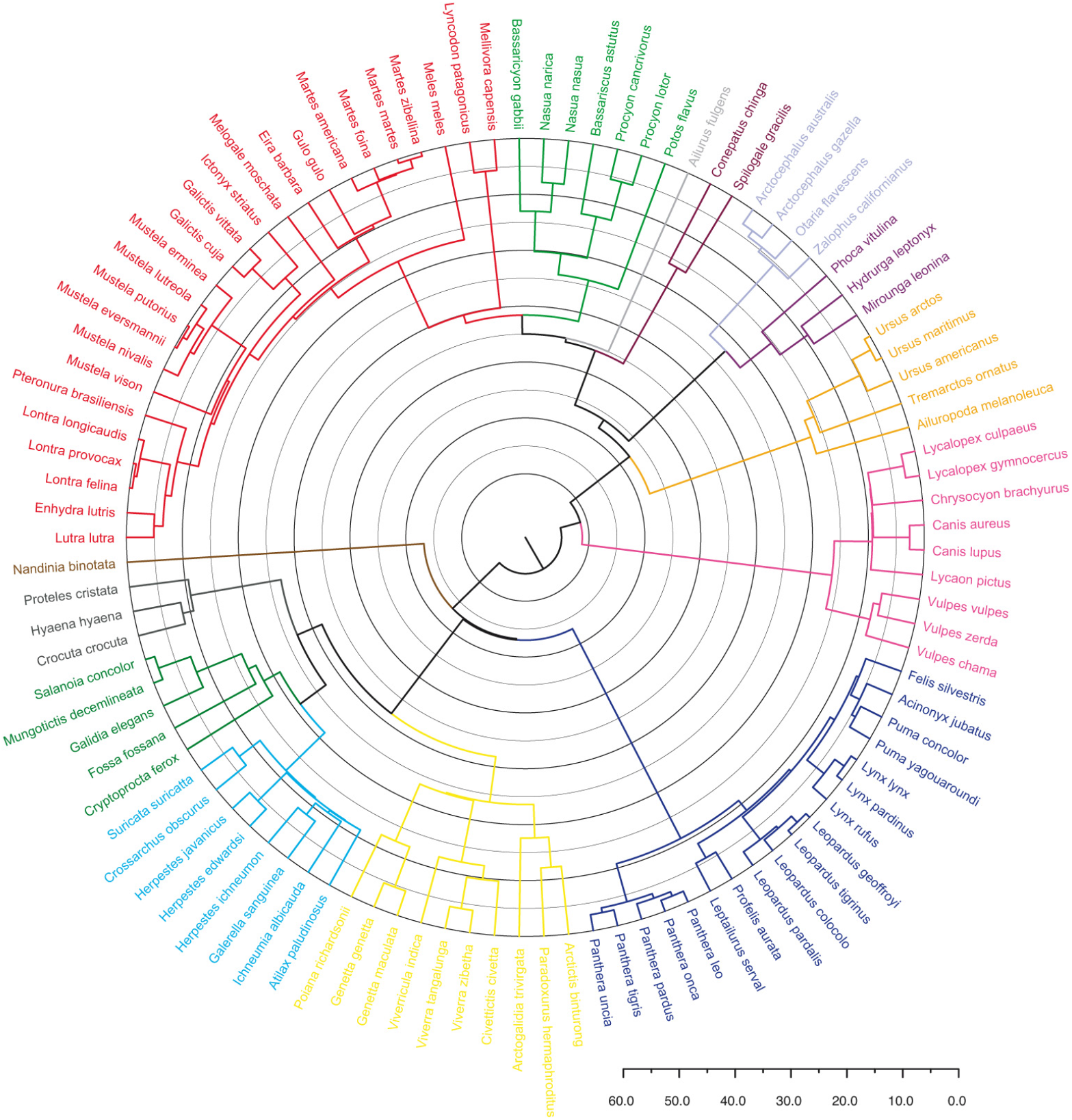
Phylogenetic relationships among the carnivoran species included in this study. The timescale represents divergence times in millions of years. The phylogeny shown was modified after Nyakatura & Bininda-Emonds (2012), as described in the Supplementary Materials of Gálvez-López (2021).

Specimens studied are housed in the collections of the Museu de Ciències Naturals de la Ciutadella (Barcelona, Spain), the Muséum National d’Histoire Naturelle (Paris, France), the Museo Nacional de Ciencias Naturales (Madrid, Spain), the Museo Argentino de Ciencias Naturales “Bernadino Rivadavia” (Buenos Aires, Argentina), and the Museo de La Plata (La Plata, Argentina). Only adult specimens (judged by epiphyseal fusion in the forelimb long bones) were sampled and, where possible, only the left scapula was digitized.

To describe scapula shape, the three-dimensional coordinates of 34 landmarks (15 true landmarks and 17 semilandmarks) were recorded using a Microscribe G2X digitizer (Immersion Corporation; San Jose, California, US) (Fig. 2; Table 3). After digitalization, semilandmark coordinates were recalculated with RESAMPLE (Raaum, 2006), which uses weighted linear interpolation to evenly space semilandmarks along the curve they define.

**Figure 2.**
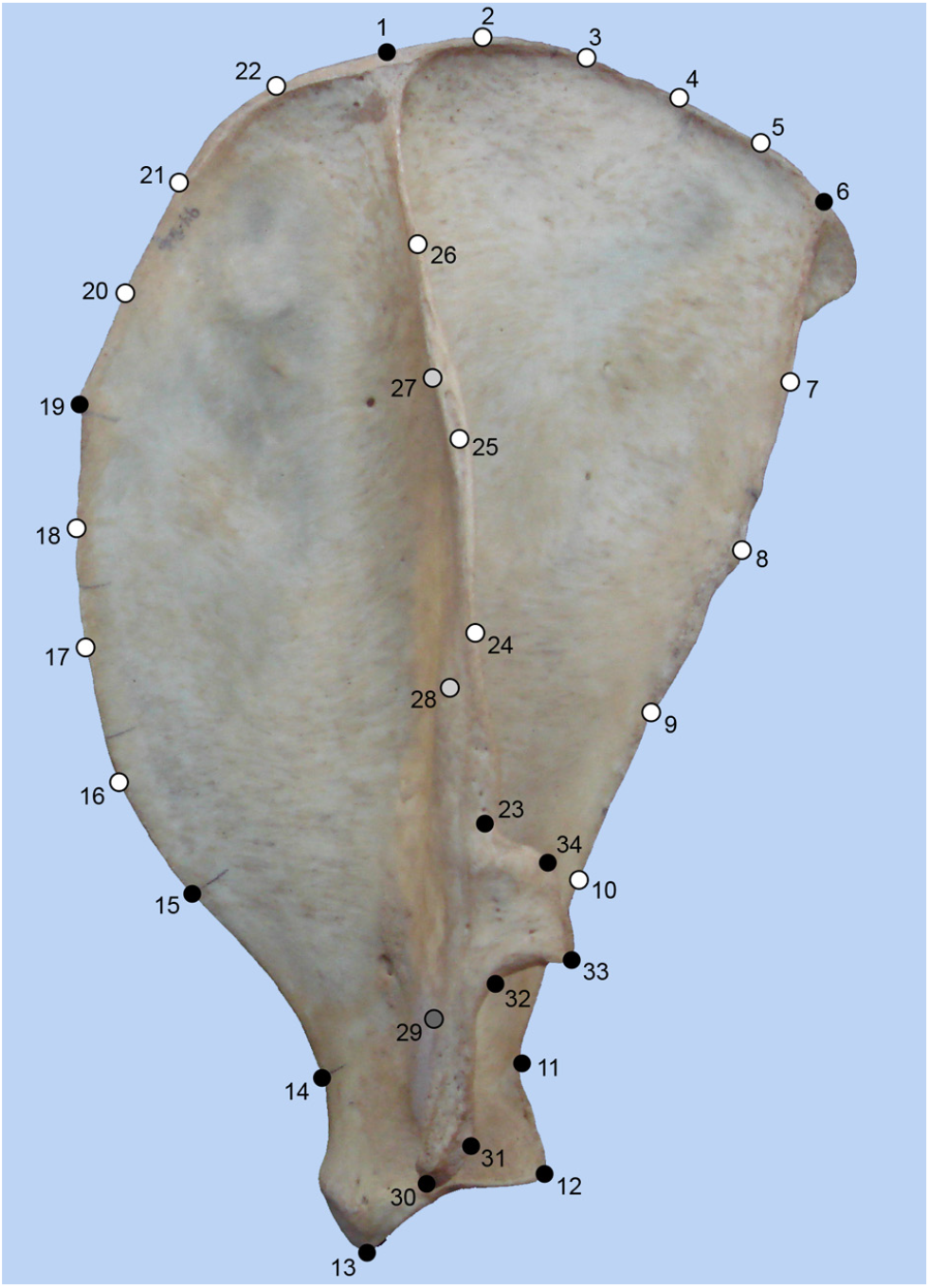
Left scapula of *Acinonyx jubatus* showing the landmark configuration used in this study. Black dots represent true landmarks, while white dots correspond to semilandmarks. Landmark 29 and semilandmarks 27 and 28 are greyed out because their anatomical position cannot be observed directly on the picture. See Table 3 for definition of landmarks.

**Table 3.** Scapular landmarks used in this study.

| Landmark | Definition |
| --- | --- |
| 1 | Intersection between scapular spine and vertebral border. |
| 2 – 5 | Semilandmarks along caudal part of vertebral border. |
| 6 | Caudal angle. |
| 7 – 10 | Semilandmarks along caudal border. |
| 11 | Point of maximum curvature along caudal scapular neck margin. |
| 12 | Caudalmost point on lip of glenoid cavity. |
| 13 | Cranialmost point on lip of glenoid cavity. |
| 14 | Point of maximum curvature along cranial scapular neck margin. |
| 15 | Ventral end of cranial border. |
| 16 – 18 | Semilandmarks along cranial border. |
| 19 | Cranial angle. |
| 20 – 22 | Semilandmarks along cranial part of vertebral border. |
| 23 | Limit between acromion and scapular spine. |
| 24 – 26 | Semilandmarks along edge of scapular spine. |
| 27 – 28 | Semilandmarks along base of scapular spine. |
| 29 | Ventralmost point of scapular spine base. |
| 30 | Cranialmost point of tip of acromion. |
| 31 | Caudalmost point of tip of acromion. |
| 32 | Point of maximum curvature between acromion and metacromion. |
| 33 | Ventralmost point of tip of metacromion. |
| 34 | Dorsalmost point of tip of metacromion. |

Measurement error (ME) was quantified using partial superimposition (von Cramon-Taubadel et al., 2007). For that, the landmark configurations of five specimens belonging to five different-sized species were digitized five times (Aju, Pco, Plo, Mfo, Vvu; see Table 1 for species names abbreviations). Landmarks 1, 6 and 29 were used as the baseline. The mean measurement error for any given landmark ranged between 0.24 and 0.48 mm (i.e., close to once and twice the accuracy of the digitizer, ± 0.23 mm, respectively). The highest ME values corresponded to landmarks 18 and 19, which reflected the difficulty of precisely locating the cranial angle in some species.

Prior to any further analyses, all non-shape information (i.e., size, location and orientation) was removed performing a Generalized Procrustes Superimposition on all landmark configurations. Briefly, this procedure first standardizes size by equaling centroid size (CS; the squared root of the sum of the squared distances of each landmark to the centroid of the configuration) to unit size in each configuration, then shifts all configurations so that their centroid is located at the same position, and finally aligns the configurations by iteratively minimizing the sum of squared distances between corresponding landmarks of each configuration. The scaling of CS to unit size only removes isometric size effects, retaining both shape variation unrelated to size and allometric shape variation. A more detailed explanation of this procedure and its computation can be found elsewhere (Bookstein, 1991; Dryden & Mardia, 1998; Zelditch et al., 2004). After the Procrustes superimposition, the aligned landmark configurations (Procrustes coordinates) were averaged by species to eliminate the possible effect of static allometry and sexual shape dimorphism.

The allometric effect on scapula shape was quantified by regressing the Procrustes coordinates (shape variables) onto CS. Due to the large body size range in the species sampled, the regression was performed using both raw CS data and log-transformed values (log CS). Since the latter magnifies shape changes at small sizes, differences between both regressions would suggest that the allometric effect changes along the studied size range. This is known as differential scaling and it has been previously reported for several linear measurements in carnivoran limb bones (Bertram & Biewener, 1990; Gálvez-López & Casinos, 2022*a*, *b*).

The presence of a phylogenetic signal in the shape of the carnivoran scapula was tested using a permutation test (Klingenberg & Gidaszewski, 2010). In this test, the null hypothesis of complete absence of a phylogenetic signal is simulated by randomly permuting the Procrustes coordinates among the terminal taxa and calculating the total amount of squared change summed over all branches of the tree. Then, these summed squared changes are compared to the value calculated for the original data, and an empirical p-value is thus defined as the proportion of permutated data sets with sums lower or equal than the original.

In addition, a MANOVA by locomotor type was carried out on the Procrustes coordinates to determine whether locomotor differences had a significant effect on scapula shape in Carnivora.

Shape variation in the carnivoran scapula was first explored using a principal components analysis (PCA) of the shape variables. The number of principal components (PCs) to be further analyzed was determined using the broken-stick model (Frontier, 1976), according to which a PC can be interpreted if its observed eigenvalue exceeds the value expected under a random distribution of total variance amongst all PCs. Then, the effect of size, phylogeny and function (locomotor type) was evaluated in each of those PCs individually. The former was quantified regressing the PC scores onto CS (and log CS), while one-way Procrustes ANOVAs were used to assess the effect of the other two factors. Additionally, the reconstructed ancestral shapes (see below) were plotted onto the shape spaces defined by the PCs and then connected by the branches of the tree in Figure 1. The resulting phylomorphospaces allow the assessment of the evolutionary history of shape changes (Klingenberg & Gidaszewski, 2010).

While PCA identifies the major axes of shape variation, it is not intended to separate those specimens into groups. Thus, a canonical variates analysis (CVA) was performed on the shape variables to determine whether species with similar locomotor habits shared similarly shaped scapulae. Furthermore, this would allow the identification of morphological indicators for each locomotor type with a distinct scapular shape. As with PCA, the CV scores were regressed onto CS (and log CS) to quantify the effect of size on group separation. Since group means were the main target of this comparison, differences between groups should be tested based on the Mahalanobis distances between them (Klingenberg & Monteiro, 2005). However, the presence of anisotropic shape variation (results not shown), together with unequal group sizes, could violate the assumption of identical within-group covariation matrixes. Thus, discriminant function analyses (DFAs) were preferred for the intergroup comparison, and its significance was determined using permutation tests based on Procrustes distances. Furthermore, since DFA tends to over-estimate differences between groups when sample size is small compared to the number of dimensions (i.e., landmarks), the reliability of the discrimination was assessed using leave-one-out cross-validation (Lachenbruch, 1967).

Finally, hypothetical scapular sizes and shapes were reconstructed for each node of the phylogeny using squared-change parsimony weighted by branch lengths (Maddison, 1991). This procedure minimizes the sum of squared-changes along the branches of a phylogeny to reconstruct the value of continuous characters at ancestral nodes. Changes on longer branches are considered less costly because they are weighted using branch lengths (Maddison, 1991). Although this methodology has been criticized for producing wide confidence intervals for the reconstructed values, its accuracy can be increased by including fossil taxa or increasing taxon sampling (Finarelli & Flynn, 2006). Here, the latter was preferred, as complete scapulae are scarce in the fossil record due to its thinness. Once the ancestral scapular sizes and shapes were reconstructed, the scapular morphology of living species was used to infer locomotor type at several nodes.

All analyses were performed using the software package MorphoJ (Klingenberg, 2011), except for the ANOVAs and MANOVAs, which were carried out with SPSS for Windows (release 15.0.1 2006; SPSS Inc., Chicago, IL, USA).

## Results

The shape variation of the carnivoran scapula is significantly influenced by both size and phylogeny. The regressions of the shape variables (Procrustes coordinates) onto CS and log CS produced similar results, indicating that about 17% of scapular shape variation is caused by size differences among species (CS: 16.71% / log CS: 17.18%; p < 0.0001 in both cases) (Fig. S1). Furthermore, the permutation test revealed a strong phylogenetic signal in scapular shape (p < 0.0001). Together with the MANOVA performed on the Procrustes coordinates, which revealed significant shape differences between locomotor types (p < 0.0001), these results indicate that scapular morphology responds to the complex interaction of allometric, phylogenetic, and functional effects.

### Shape variability

The broken-stick model suggested that the first eleven PCs could be interpreted. Table 4 shows the results for the allometric regressions and the ANOVAs by family and locomotor type for each of those PCs. Each of the first four PCs was significantly related to size, phylogeny and locomotor type (Table 4). On the other hand, PC5 through PC11 showed a varied relationship with these factors (Table 4), being significantly related either to one, two, or none of them.

**Table 4.**
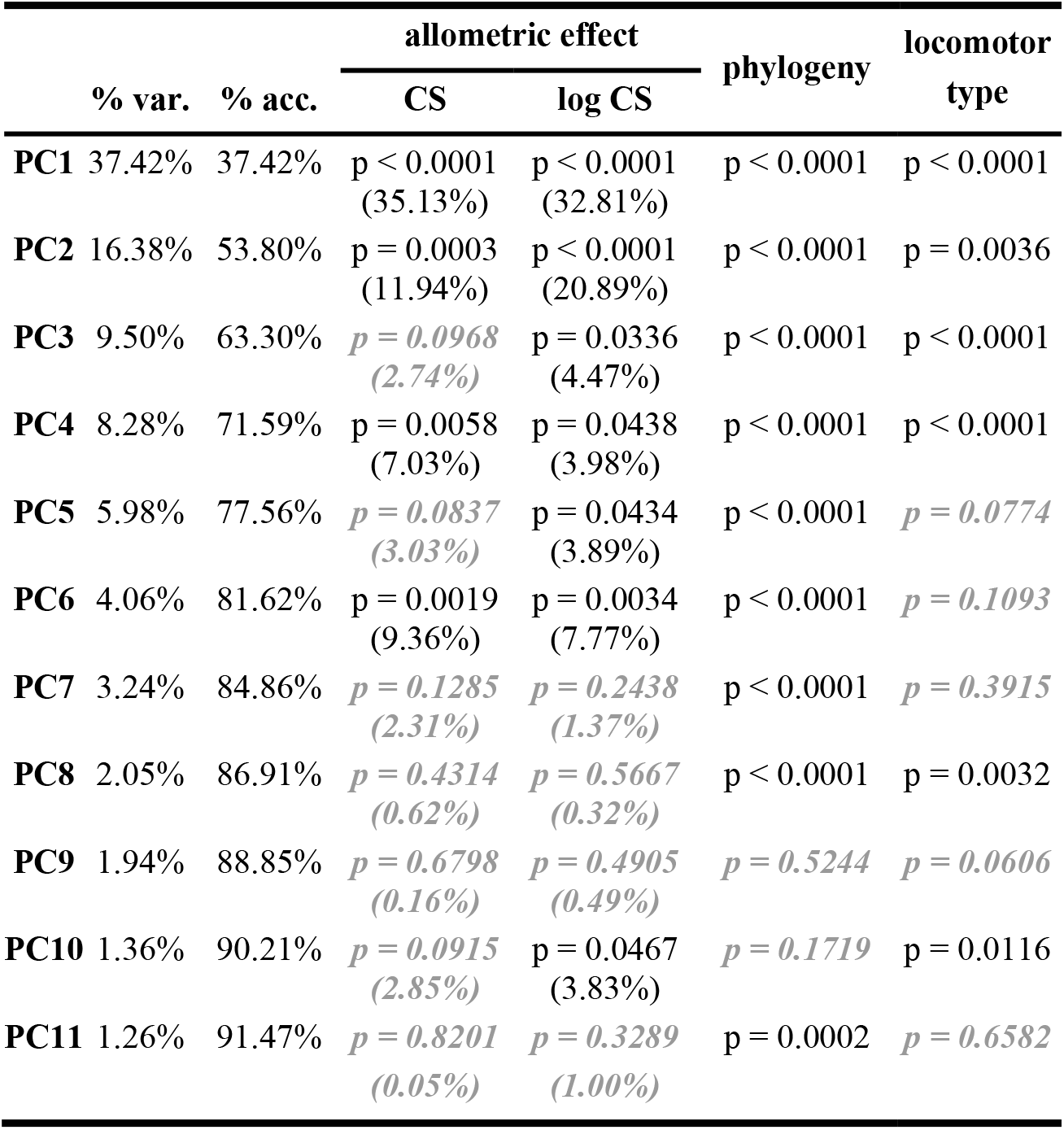
Effect of size, phylogeny and locomotor type in the first principal components. The allometric effect on each principal component (PC) was determined using regression methods, which allows the shape variation explained by size changes to be expressed as a percentage of the total (values in parentheses). Non-significant results (i.e., p-value > 0.05) are presented in ***grey bold italics***. Abbreviations: % acc., accumulated percentage of explained shape variance; % var., percentage of shape variance explained by eachPC; CS, centroid size.

PC1 explained 37.42% of the shape variation of the carnivoran scapula, clearly separating pinnipeds and fissipeds (Fig. 3). Extreme negative PC1 values corresponded to short and wide scapulae with large fossae, a broad neck, and an almost nonexistent acromion. On the other hand, extreme positive PC1 values were associated with long and narrow scapulae with a high spine, well-developed acromial processes (hamatus and suprahamatus), and an extremely reduced anterior part of the vertebral border.

**Figure 3.**
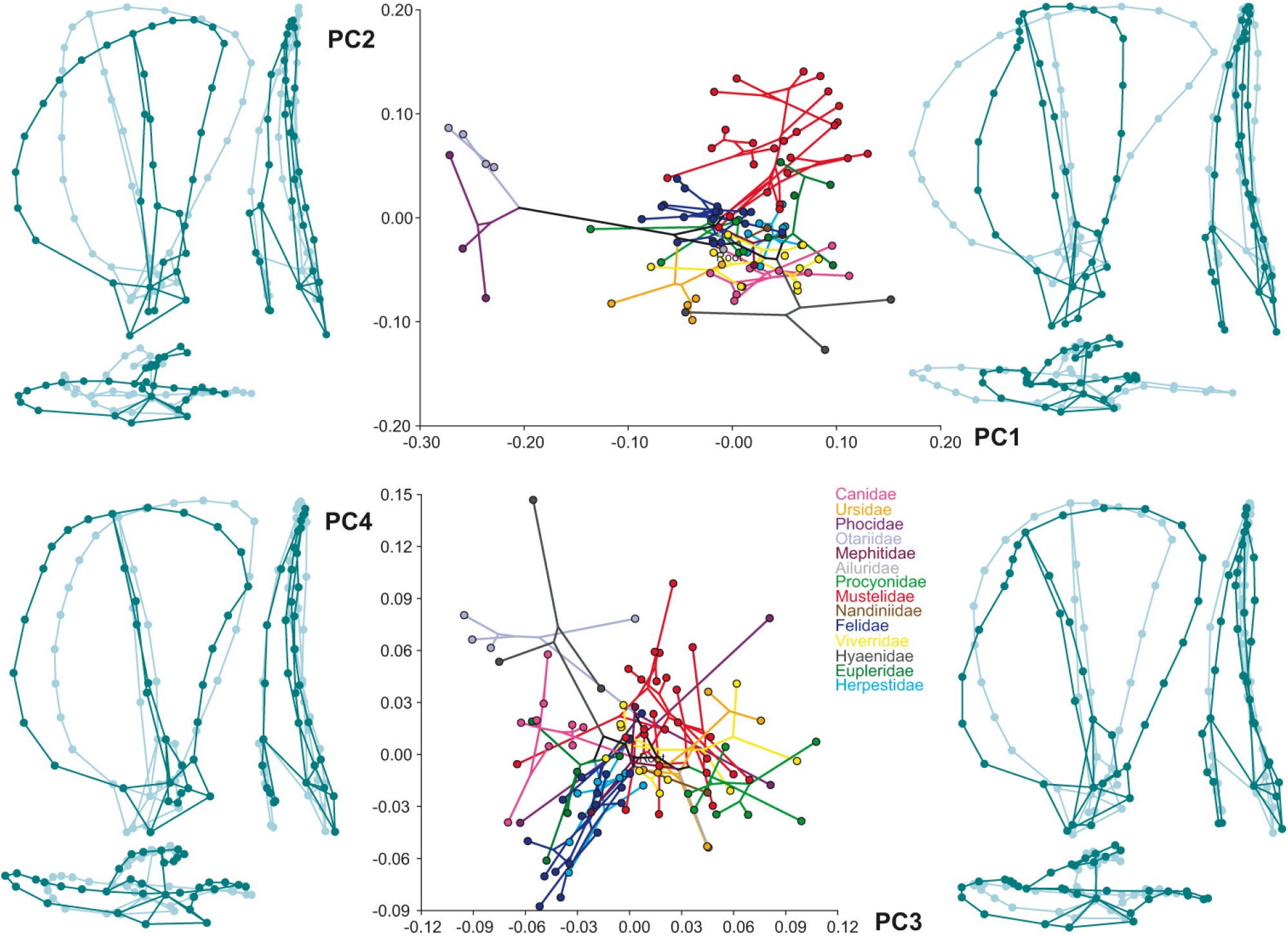
Principal components analysis of shape variation in the carnivoran scapula. The plots represent the phylomorphospaces defined by the first four principal components (PCs), which explained over 70% of shape variation. The tree topology projected on each phylomorphospace corresponds to the phylogeny presented in Figure 1. The shape changes associated to each PC are displayed using wire-frames, from the most negative values (light blue) to the most positive ones (dark blue). For each PC, a set of three pairs of wire-frames is presented so that shape changes can be observed in lateral (x-y; left), dorsal (x-z; bottom), and caudal (y-z; right) views.

The shape changes associated to PC2 (16.38%) were mainly related to the relative development of the fossae. Species with the lowest PC2 values presented an expanded infraspinous fossa and a smaller supraspinous fossa, while the opposite was true for those with high PC2 values (Fig. 3). This expansion of the supraspinous fossa along PC2 was accompanied by its flattening. Additionally, with increasing PC2 values, the dorsal end of the spine displaced caudally as the cranial part of the vertebral border expanded and the caudal part contracted. Finally, as in PC1, the acromion processes were poorly developed at the negative end and clearly distinct at the positive end.

Together, PC3 and PC4 accounted for 17.78% of shape variation (Table 4). Increasing PC3 values resulted in the expansion of the caudal part of the vertebral border and the contraction of the cranial part, which resulted in the cranial displacement of the dorsal spine (Fig. 3). Furthermore, both the cranial and caudal borders expanded, increasing scapular width; while the well-developed acromion became larger and wider, and twisted its orientation relative to the scapular blade. Regarding PC4, the cranial border expanded with increasing PC4 values, reducing the cranial part of the vertebral border and enlarging the supraspinous fossa (Fig. 3). On the other hand, the caudal part of the vertebral border shifted distally at the caudal angle, while the glenoid cavity displaced caudally, which resulted in a short and concave caudal border. Furthermore, the acromion processes shrank and the scapular blade increased its medio-lateral curvature.

Other than the separation of pinnipeds, no clear phylogenetic pattern could be observed in the phylomorphospace defined by those pairs of PCs (Fig. 3). Similarly, no clustering of locomotor types became apparent using those axes (Fig. S2). This is probably related to those PCs being significantly affected by size, phylogeny and function at the same time, and again suggests that scapular morphology in Carnivora results from the complex interaction of those factors.

### Allometric effect

Although the regressions of shape onto CS and log CS produced similar results, several observations could be made from the scatter plots (Fig. S1). First, pinnipeds were placed well above the main regression line (i.e., present higher shape values than similar-sized fissipeds), which could be affecting the regression results. Indeed, repeating the regressions on a subsample excluding Pinnipedia (i.e., fissiped subsample) revealed that the allometric effect decreased to 11.78% and 14.40% when using CS and log CS, respectively. Second, the allometric effect seemed to vary in the different carnivoran families (see 95% confidence ellipses in Fig. S1). And third, log-transforming centroid size did not linearize the relationship of shape and size, which in bivariate scaling studies suggests the presence of differential scaling (Bertram & Biewener, 1990; Gálvez-López & Casinos, 2022*a*, *b*).

Regarding the allometric effect on the different PCs, the regressions on CS and log CS produced different result in several cases. In the case of PC2 size explained about 12% of shape variation when using raw CS data, but close to 21% when using log CS. Similarly, the regression on log CS for PC3, PC5 and PC10 were significant but not those using raw CS data (Table 4). This would suggest that the shape changes described by those PCs are more strongly affected by size in small species than in large carnivorans. On the other hand, the shape changes described by PC4 should be more accentuated in large species, since the magnitude of the allometric effect almost halves and is barely significant when using log CS (Table 4).

In light of these results, the effect of size on scapular shape was further explored, separately, in the different carnivoran families and locomotor types studied (Table S1). Overall, the regressions on CS and log CS produced similar results, which suggested that the differential scaling observed in scapular shape is a consequence of scaling differences among carnivoran families and/or among locomotor types, not among different-sized species. For instance, the allometric effect detected in Felidae, Mustelidae, and scansorial carnivorans, was consistent, no matter the size variable used, despite the wide size ranges of those groups. A significant allometric effect was found in Mustelidae, Felidae, and Herpestidae, and in scansorial, terrestrial, semifossorial, and semiaquatic carnivorans (Table S1). It is interesting to note that, although the amount of shape variation explained by size differences was large in Ursidae, Procyonidae, and Eupleridae, the regressions were not significant, probably due to the small sample size of those families.

In order to further explore the relationship between shape, size and phylogeny, the regression of shape variables on log CS was repeated using phylogenetically independent contrasts (PIC). This methodology incorporates the phylogenetic structure of the sampled species into the analysis, and thus accounts for the potential correlation of the error terms that could arise due to the lack of independence among species (Felsenstein, 1985). The allometric effect was still significant when taking into account the phylogenetic signal in the studied dataset, but the amount of shape variation explained by size dropped to 5.31% (5.75% in the fissiped subsample). These results suggest that most of the allometric shape differences found in the carnivoran scapula are related to size changes along phyletic lines, and supports that differential scaling in scapular shape is caused by the juxtaposition of different family-specific allometric trajectories. Finally, when studying the effect of size on scapular shape in the different families and locomotor types, PIC regressions only showed a significant allometric effect in Mustelidae, Herpestidae, and scansorial carnivorans (Table S1). Furthermore, the amount of scapular shape variation explained by size was greatly reduced in all subsamples but herpestids. These results also indicate that, in the rest of subsamples for which a significant allometric effect was reported above, allometric shape differences were related exclusively to size changes along phyletic lines.

Summarizing, the results of the present study suggest that there is evidence for differential scaling in the shape of the carnivoran scapula, which is mainly caused by scaling differences among carnivoran families. In fact, although size explains about 17% of scapular shape variation in Carnivora, only 5.3% of this allometric shape variation is not related to size changes along phyletic lines.

### Shape and locomotion

The CVA on shape variables grouping species by locomotor type produced six non-zero CVs. The first two accounted for 82.37% of shape variation (Table 5) and defined a shape space where all groups were clearly separated (Fig. 4). However, pairwise DFAs could not significantly separate semifossorial from semiarboreal species, nor scansorial from terrestrial carnivorans, nor semiarboreal from terrestrial species (Table 6). Small sample sizes probably affected the resolution of DFAs, but the rather broad definition of the terrestrial category might have collaborated as well.

**Table 5.** Canonical variates analysis of scapular shape by locomotor type. The allometric effect on each canonical variate (CV) was determined using regression methods, which allows the shape variation explained by size changes to be expressed as a percentage of the total (in parentheses). Non-significant results (i.e., p-value > 0.05) are presented in ***grey bold italics***. Abbreviations: % acc., accumulated percentage of explained shape variance; % var., percentage of shape variance explained by each CV; CS, centroid size.

|  | % var. | % acc. | allometric effect |  |
| --- | --- | --- | --- | --- |
|  |  |  | CS | log CS |
| CV1 | 63.58% | 63.58% | p < 0.0001<br>(23.30%) | p < 0.0001<br>(17.67%) |
| CV2 | 18.79% | 82.37% | *p = 0.6808*<br>*(0.17%)* | *p = 0.4004*<br>*(0.70%)* |
| CV3 | 11.94% | 94.31% | *p = 0.1804*<br>*(1.82%)* | *p = 0.1813*<br>*(1.78%)* |
| CV4 | 2.98% | 97.29% | *p = 0.5060*<br>*(0.48%)* | *p = 0.5352*<br>*(0.39%)* |
| CV5 | 1.88% | 99.16% | *p = 0.6879*<br>*(0.17%)* | *p = 0.5420*<br>*(0.37%)* |
| CV6 | 0.84% | 100.0% | p = 0.0021<br>(8.70%) | p = 0.0007<br>(10.87%) |

**Figure 4.**
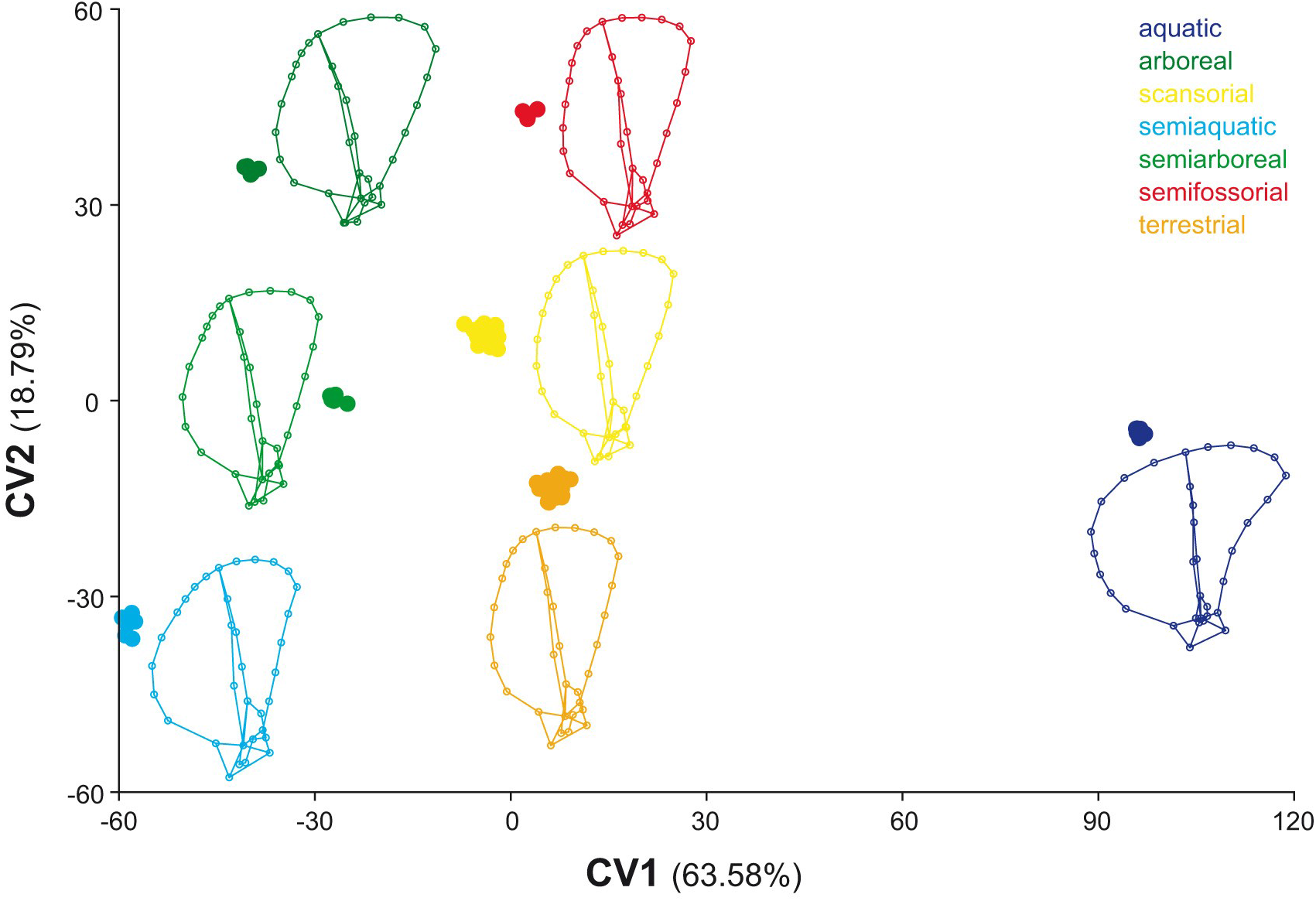
Canonical variates analysis of shape variation in the carnivoran scapula, grouped by locomotor type. Percentages express the amount of shape variance explained by each canonical variate (CV). The mean shape of each locomotor type categories is also presented (in lateral view).

**Table 6.**
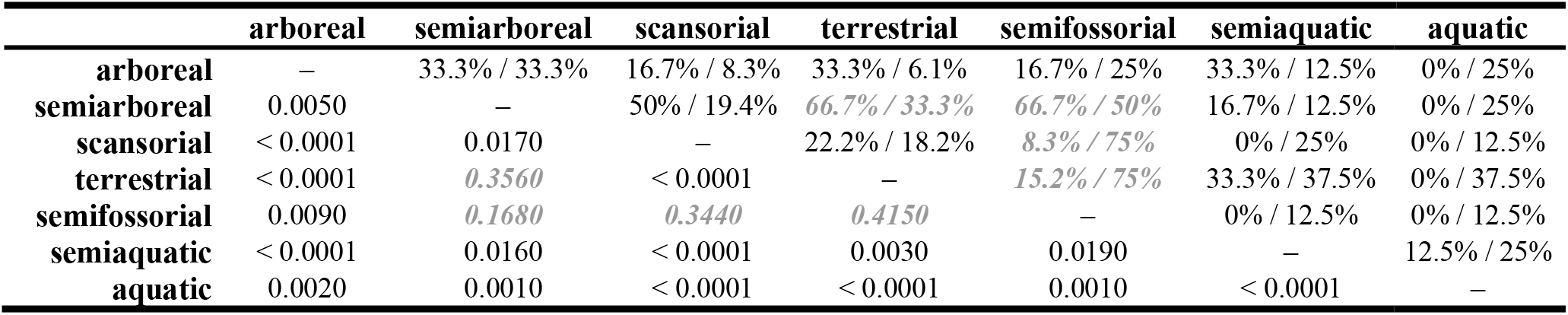
Shape differences between locomotor types. Each pair of locomotor types was compared using discriminant function analysis based on Procrustes distances. The p-value for each of the pairwise comparisons is shown under the diagonal, whereas values over that line represent misclassification rates of the cross-validation procedure. The first percentage indicates the amount of row-group species misclassified as column-group carnivorans, while the second percentage corresponds to column-groups species incorrectly placed into the row-group. Results of non-significant pairwise comparisons are presented in ***grey bold italics***.

|  | arboreal | semiarboreal | scansorial | terrestrial | semifossorial | semiaquatic | aquatic |
| --- | --- | --- | --- | --- | --- | --- | --- |
| arboreal | — | 33.3% / 33.3% | 16.7% / 8.3% | 33.3% / 6.1% | 16.7% / 25% | 33.3% / 12.5% | 0% / 25% |
| semiarboreal | 0.0050 | — | 50% / 19.4% | *66.7% / 33.3%* | *66.7% / 50%* | 16.7% / 12.5% | 0% / 25% |
| scansorial | < 0.0001 | 0.0170 | — | 22.2% / 18.2% | *8.3% / 75%* | 0% / 25% | 0% / 12.5% |
| terrestrial | < 0.0001 | *0.3560* | < 0.0001 | — | *15.2% / 75%* | 33.3% / 37.5% | 0% / 37.5% |
| semifossorial | 0.0090 | *0.1680* | *0.3440* | *0.4150* | — | 0% / 12.5% | 0% / 12.5% |
| semiaquatic | < 0.0001 | 0.0160 | < 0.0001 | 0.0030 | 0.0190 | — | 12.5% / 25% |
| aquatic | 0.0020 | 0.0010 | < 0.0001 | < 0.0001 | 0.0010 | < 0.0001 | — |

Since the previous analyses suggested a strong interaction between phylogeny and size in scapular shape (see above), the CVA and DFAs were repeated on the residuals of the PIC regression of shape variables onto log CS. The results of these size- and phylogeny-corrected analyses were practically identical to those of the original dataset (results not shown), suggesting that locomotor indicators in scapular shape are independent of size or shared ancestry in Carnivora. Thus, only the results of the uncorrected analyses are further discussed, since they provide a more realistic comparison for reconstructed ancestral shapes.

The first CV explained 63.58% of scapular shape variation and was significantly affected by size (Table 5). CV1 clearly separated aquatic carnivorans on one extreme and semiaquatic species on the other, while the remaining locomotor type categories occupied an intermediate position with increasing CV1 scores as arboreality decreased (Fig. 4). The shape changes associated with increasing CV1 scores corresponded to an expansion of the vertebral border and a flattening of the scapular blade, coupled with a reduction and reorientation of the acromion processes. Furthermore, the scapular neck narrowed and the spine straightened from a cranially bent position.

The highest and lowest CV2 scores corresponded, respectively, to semifossorial and semiaquatic carnivorans (Fig. 4). Increasing CV2 scores described a reorientation of the glenoid cavity and the acromion, the former aligning with the scapular blade from a medially oriented position and the latter shifting craniolaterally. Furthermore, the infraspinous fossa expanded and the supraspinous fossa slightly contracted.

### Ancestral state reconstruction

Reconstructed ancestral states for scapular size and shape are presented, respectively, in Figures 5 and 6. The basal carnivoran had a centroid size of 247.92 mm, which corresponds to the scapula size of some extant medium-sized species (e.g. most lynxes (*Lynx sp*.), the larger otters (*Enhydra, Pteronura*), and the aardwolf (*Proteles*)). In caniform carnivorans, scapula size first augmented, leading to increasingly larger clades (Canidae, Ursidae, Pinnipedia), and then decreased in the musteloid clades (Fig. 5). On the other hand, scapula size decreased during the evolution of most feliform clades, increasing only in Felidae and Hyaenidae (Fig. 5). The scapula shape reconstructed for the carnivoran ancestor did not resemble any of the shapes of the extant species (Fig. 6A). The shape of its scapular blade roughly corresponded to that of some genets and mongooses, but in the former both the insertion of m. levator scapulae and the acromion processes were shorter, while in the latter the vertebral border extended more dorsally and both spine and acromion had a different shape. However, this ancestral shape closely matched the mean shape of scansorial carnivorans (Figs. 4, 6A), which suggests that the carnivoran ancestor was a good climber but spent most of its time on the ground. The main differences between the ancestral shape and the scansorial mean were a slightly larger infraspinous fossa and an acromion similar to the mean terrestrial shape, although more cranially oriented (as in scansorial species) (Figs. 4, 6A). The former differences indicate a higher degree of arboreality in this ancestor than in extant scansorial carnivorans, while the latter could be interpreted as a different solution for a similar problem (i.e., a moderate degree of arboreality).

**Figure 5.**
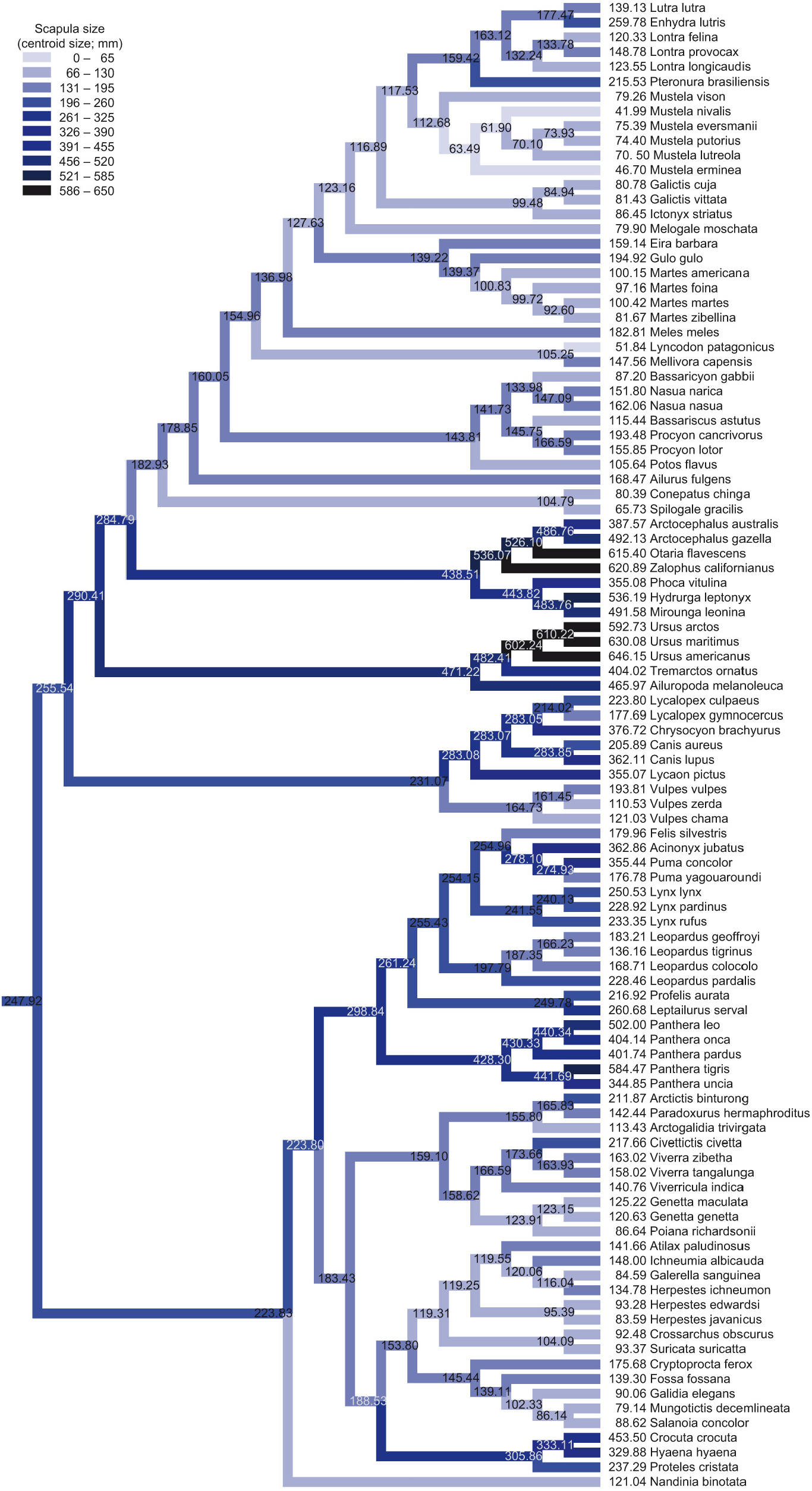
Ancestral states reconstruction of scapular size. Values at terminal nodes represent mean centroid size (CS) of the scapula for each species studied, while values at internal nodes correspond to estimated ancestral sizes. Branch shading corresponds to estimated CS values and is maintained in Figure 6 for cross-reference. Note that branch lengths are not drawn at their proportional length to ease the visualization of the results (see Fig. 1 for actual branch lengths).

**Figure 6.**
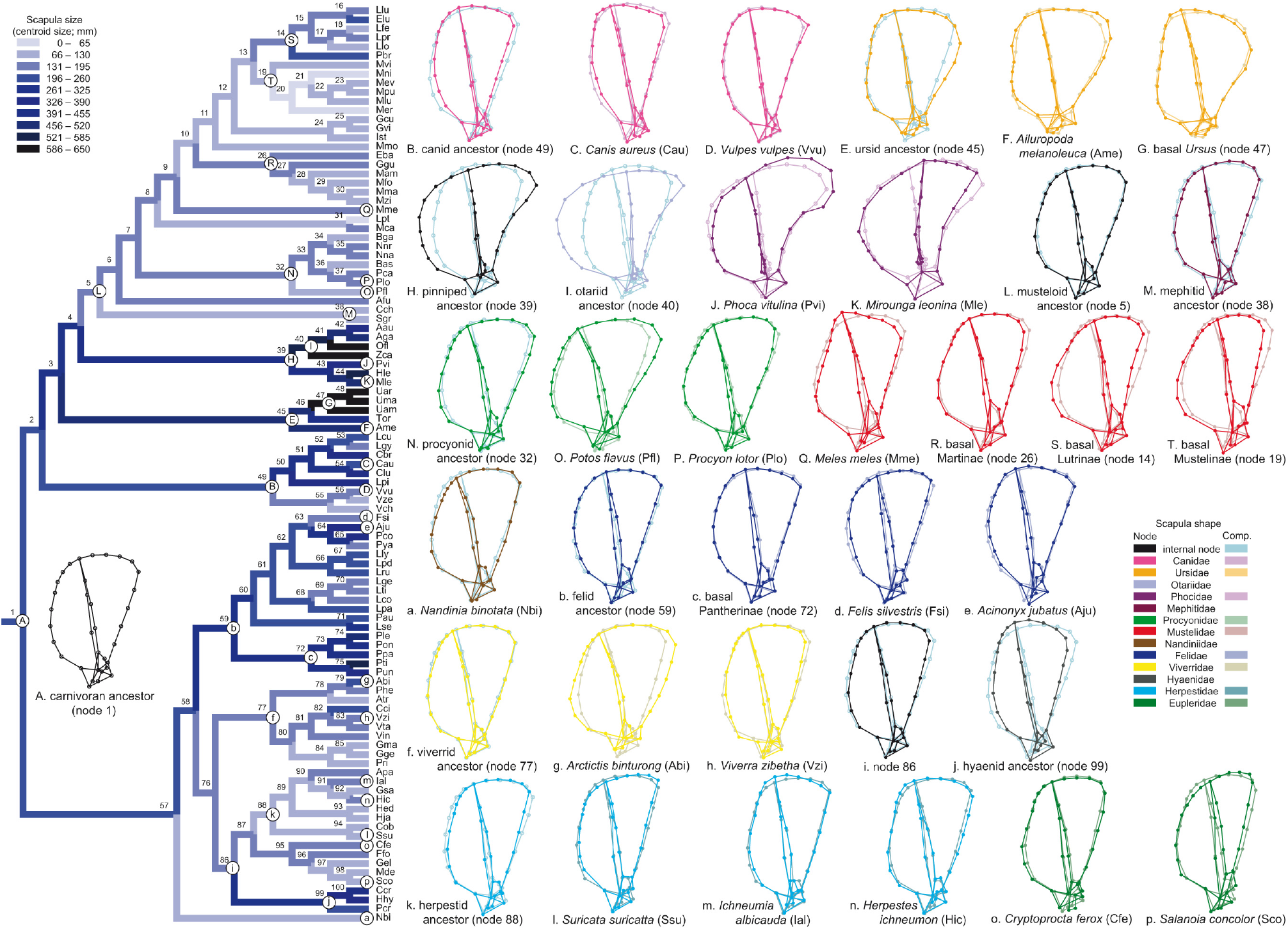
Ancestral states reconstruction of scapular shape. Wire-frames correspond to either selected estimated internal nodes or to mean scapular shapes of extant species. For simplicity, only the most relevant nodes are shown. The shape of the carnivoran ancestor (A) and those of Caniformia are labeled with upper case letters (B, C,…), while Feliformia uses lower case letters (a, b,…). Internal nodes common to several families (H, L, i), plus the ancestor of each family (e.g. B, b), are compared to the estimated shape of the carnivoran ancestor (node A). Terminal taxa (e.g. C, d) and internal nodes within families (e.g. G, c) are compared to the ancestor of their family instead (e.g. C to B). All scapular shapes are presented in lateral view. Branch shading in the phylogetic tree corresponds to estimated CS values (Fig. 5). Note that branch lengths are not drawn at their proportional length to ease the visualization of the results (see Fig. 1 for actual branch lengths). *Legend:* “Node” denotes the main color used on the wire-frames of each family, while “Comp.” is the color used on the comparative wire-frames. See Table 1 for species names abbreviations.

From this ancestral shape, the carnivoran scapula suffered several major shape changes along the branches leading to the extant families. Thus, the evolution of the carnivoran scapula is more deeply addressed in the discussion of these results.

## Discussion

### Morphological variation in the carnivoran scapula

According to previous studies, the shape of the mammalian scapula is strongly linked to shared evolutionary history and, within each lineage, shape variability can be attributed either to size differences between closely related species (Didelphidae, Astúa, 2009), or to different functional requirements (Xenarthra, Monteiro & Abe, 1999; Caviomorpha, Morgan, 2009; Anthropoidea, Young, 2008). However, the results of the present study suggest that, in Carnivora, scapula shape reflects the complex interaction of historical, allometric and functional effects. Not only can none of these factors alone explain the large shape variability of the carnivoran scapula (Figs. 3, S1, S2), but also this shape variability is highly correlated to all of those factors, considering either scapula shape as a whole (Procrustes coordinates, see Results) or the main axes of shape variation separately (PC1 to PC4, Table 4). Previous studies on forelimb bones have reported a similar strong interaction between size, phylogeny and function in Carnivora (Walmsley et al., 2012; Fabre et al., 2013*a*).

Besides defining the main axes of shape variation, the PCA results also identified the scapular regions in which most of this variation occurred. That is, while shape changes in most scapular regions were described by one or few PCs (e.g. PC2 and PC4 defined the angle between the spine and the glenoid cavity), those of some other regions were described by most PCs, suggesting that they represent most of the shape variability of the carnivoran scapula. One of these regions was the vertebral border, in which the serratus ventralis muscle (= m. serratus anterior *sensu lato*) inserts. This muscle can be subdivided into an anterior part (m. levator scapulae), which inserts on the cranial part of the vertebral border, and a posterior part (m. serratus anterior *sensu stricto*), inserting on the caudal part of the vertebral border. The former protracts the forelimb, while the latter retracts it (Maynard Smith & Savage, 1956; Argot, 2001). The major shape changes observed in the vertebral border were its overall expansion/contraction, the relative length of its cranial and caudal parts, and the angulation between these parts (Fig. 3). All of these features determine the size and moment arms of mm. levator scapulae and serratus anterior *s. s*., and have already been related to several locomotor specializations (Oxnard, 1968; Taylor, 1974; English, 1977; Argot, 2001; Astúa, 2009; Morgan, 2009). A second region of high shape variability were the fossae, which varied mainly in their overall extension and in their relative development (i.e., whether the supraspinous or infraspinous fossa was larger) (e. g. PC1–PC3; Fig. 3). Larger scapular fossae reflect enlarged supraspinous and infraspinous muscles (Roberts, 1974). Besides its main function as shoulder stabilizers, these muscles play an important role in shoulder mobility (Argot, 2001): m. supraspinatus protracts the humerus, while m. infraspinatus rotates it. Furthermore, both support humeral abduction. Thus, the relative development of the fossae indicates whether greater forelimb protraction or rotation is required during locomotion. Finally, another common feature of most PCs was the degree of development of the acromial processes (e.g. PC1, PC2, PC4; Fig. 3). The hamatus process provides insertion to the acromiodeltoid muscle, the main abductor of the forelimb, while the acromiotrapezius and omotransversarius muscles attach to the suprahamatus process and move the scapula cranially (Larson, 1993; Argot, 2001). Furthermore, the orientation of the hamatus process has been related to several locomotor adaptations (Maynard Smith & Savage, 1956; Lehmann, 1963; Roberts, 1974; Taylor, 1974; Argot, 2001).

The fact that most shape variation in the carnivoran scapula occurs in the vertebral border, fossae, and acromion, would explain the different results obtained here and in a previous study (Martín-Serra et al., 2014). In that study, the authors concluded that the main axis of shape variation in the carnivoran scapula (their PC1) corresponded to the transition “from the long and slender scapula of canids and procyonids […] to the wide and robust one of bears” (Martín-Serra et al., 2014). However, landmark selection in that study was restricted to the distal scapula (acromion, scapular neck and glenoid) plus two landmarks encompassing the origin of the teres muscle (caudal border), and thus largely omitted the major regions of shape variation characterizing scapula length and width (vertebral border, fossae). Further discrepancies between the results of the present study and those of Martín-Serra et al. (2014) were the amount of scapula shape variation explained by the allometric effect (significantly higher in the present study), and the evolutionary history of shape changes in the carnivoran scapula. While the previous study found that the different carnivoran families occupied well-differentiated portions of the morphospaces, consistent overlap was observed in the present study. Both discrepancies could be related to the different species composition of both studies. While the present study focuses on extant diversity of Carnivora, Martín-Serra et al. (2014) aimed to reconstructing the evolutionary history of large-bodied families, including both extant and extinct species. The higher percentages of allometric shape variation obtained in the present study when using log-transformed centroid size (which magnifies changes at small sizes) suggest that size has a larger effect on the shape of the scapula in small carnivorans. Thus, a lower allometric effect would be expected in a sample consisting mainly of large species, as that of Martín-Serra et al. (2014). In fact, removing from the sample the species not measured in the previous study resulted in the allometric effect explaining around 10% of scapula shape variation, very close to the value reported by Martín-Serra et al. (2014). Another possible explanation would be that the shape of the fossae and spine was more heavily influenced by size differences than that of the acromion or coracoglenoid regions. However, this explanation is unlikely, since shape variation between the scapular blade and the acromion is highly integrated (Young, 2004). Furthermore, the regression of shape on size in a subsample consisting only of landmarks measured on the scapular blade and spine produced similar results to that performed on a subsample including only acromion and glenoid landmarks (results not shown). Finally, regarding phylomorphospace occupation, if only the living species measured by Martín-Serra et al. (2014) are represented in the morphospaces shown in Figure 3, most remaining families are confined to separate regions of the morphospace, mirroring the results of the previous study. In summary, comparison of our current results and those of Martín-Serra et al. (2014) emphasize the importance of increased taxon sampling in evolutionary studies (Finarelli & Flynn, 2006).

### Differential scaling on scapula shape in Carnivora

The results of this study strongly support the presence of diferential scaling in the shape of the carnivoran scapula (i.e., the amount of shape variation explained by size differences varies with increasing size). One of the arguments favoring differential shape scaling is the higher shape values found in pinnipeds relative to similar-sized fissipeds (Fig. S1). In fact, differences in centroid size explain 19.32% of scapula shape variation in pinnipeds (21.06% using log CS), whereas the allometric effect accounts for 11.78% or 14.40% of this variation in fissipeds (CS or log CS, respectively). Since size changes along phyletic lines account for most of the allometric effect on scapula shape in Carnivora, it could be argued that the scaling differences between fissipeds and pinnipeds would have a similar explanation. However, PIC regressions of shape on size also show a higher allometric effect in pinnipeds than in fissipeds (8.69% and 5.75%, respectively), confirming the scaling differences between both groups.

Further evidence for differential shape scaling in the carnivoran scapula is provided by the different percentages obtained in fissipeds when using CS and log CS as the size variable, which suggest that the allometric effect is larger in small fissipeds (since using log CS emphasizes shape changes at small sizes). In agreement with this, Martín-Serra et al. (2014) found a lower allometric effect in the shape of the carnivoran scapula using a sample of mostly large-sized fissipeds (a result that could be replicated in the present study reducing the sample to include only the species measured in the previous study, see above). Additionally, the results obtained for the different family and locomotor type subsamples also support differential shape scaling in fissipeds, since the allometric effect decreased in the groups including progressively larger species (Herpestidae > Mustelidae > Felidae; semifossorial > semiaquatic > scansorial, terrestrial; Table S1).

Summarizing, the allometric effect on scapula shape is stronger in pinnipeds than in fissipeds and, within the latter, the effect of size is more pronounced in small species. Astúa (2009) reported a similar result in Didelphidae. However, in this group scapula shape variation was more heavily influenced by size in large species, since shape differences between locomotor habits could be observed in large didelphids but all small species had similarly shaped scapulae.

### Locomotor adaptations in the carnivoran scapula

#### Swimming

The biomechanical demands of locomotion in water are highly different from those of overland locomotion (Alexander, 2002; Biewener, 2003). Water is denser and more viscous than air, which increases drag significantly. On the other hand, gravitational forces are the main factor to overcome in overland locomotion. Thus, it was not surprising that aquatic carnivorans exhibited the most characteristic scapula shape, clearly separated from all other locomotor types by CV1 (Fig. 4). As previously described for aquatic mammals (Maynard Smith & Savage, 1956), aquatic carnivorans presented the relatively shortest and widest scapulae, with a greatly expanded vertebral border (especially its cranial portion) (Figs. 4, S3). The wide vertebral border is associated with strong mm. levator scapulae and serratus anterior *s. s*., reflecting powerful forelimb protraction and retraction. These adaptations would be expected in pectoral oscillators such as otariids, which produce forward thrust beating their enlarged foreflippers (Alexander, 2002; Pierce et al., 2011). In these animals, the powerful m. levator scapulae helps overcome water drag while protracting the forelimb during the recovery phase (upstroke), while the m. serratus anterior *s. s*. retracts the forelimb during the downstroke, pushing the water downward and generating forward thrust. However, phocids swim by pelvic oscillation, generating thrust with horizontal undulations of the spine combined with hindflipper paddling (Williams, 1989; Pierce et al., 2011). Thus, aquatic locomotion cannot explain these enlarged muscles. In phocids, these muscles are probably related to terrestial locomotion, since on land the forelimbs are used to drag the heavy body, helped by axial movements. The expansion of the vertebral border, particularly its caudal portion, has also been related to an increased attachment area for the mm. teres major and deltoideus in otariids (English, 1977). In this group, the deltoid muscle consists of a single mass originating from the vertebral border near the caudal angle, extending distally along the ridge of the scapular spine into the acromion, and inserting in the deltoid tuberosity of the humerus (English, 1977). Furthermore, the caudal border of aquatic carnivorans was concave, resulting in a particularly protruding caudal angle (Figs. 4, S3), which has previously been related to an increased moment arm of the teres major muscle (Maynard Smith & Savage, 1956; Monteiro & Abe, 1999). All these features indicate strong teres major and deltoid muscles, reflecting powerful humeral adduction and abduction, respectively. In otariids, the forelimb is abducted during the upstroke and adducted during the propulsive downstroke (English, 1977). In phocids, the forelimbs provide directional control (Pierce et al., 2011), which would also involve powerful humeral abduction and adduction, not only to orient the forelimbs, but also to resist the water drag.

In agreement with previous observations on aquatic mammals (Maynard Smith & Savage, 1956), the mean scapula shape of aquatic carnivorans was the only one in which the acromion processes were completely absent (Figs. 4, S3). This is probably related to the lack of differentiation of the m. deltoideus into the spinodeltoid and acromiodeltoid muscles in pinnipeds, the latter of which originates on the hamatus process (sometimes also on the ventral lip of the suprahamatus process; English, 1977).

Finally, as previously described for otariids (English, 1977), aquatic carnivorans presented greatly expanded and flat supraspinous and infraspinous fossae, particularly the former, and a low scapular spine. As stated above, broad fossae indicate a greater development of mm. supraspinatus and infraspinatus, and thus reflect powerful shoulder stabilization but also enhanced mobility at the glenohumeral joint (Roberts, 1974; Argot, 2001; Morgan, 2009). This increase in shoulder stabilization is probably related to the greater development of humeral abductors and adductors (i.e., mm. teres major and deltoideus, see above).

Semiaquatic carnivorans shared some scapular features with aquatic species, namely a short scapular blade, a wide and flat supraspinous fossa, a markedly convex vertebral border, and a slightly concave caudal border (Figs. 4, S3). Some of these shape features were also described in the semiaquatic marsupial *Chironectes* (Astúa, 2009). However, contrary to aquatic carnivorans, semiaquatic species presented a reduced vertebral border (particularly the cranial portion), a high scapular spine with the acutest angulation relative to the scapular blade, the smallest infraspinous fossa, and well-developed acromion processes (especially the suprahamatus process) (Figs. 4, S3). The reduced insertion for m. levator scapulae would suggest weak forelimb protraction in semiaquatic carnivorans, but the action of this muscle is probably supported by the enlarged m. supraspinatus and mm. acromiotrapezius and omotransversarius (indicated, respectively, by the wide supraspinous fossa and the enlarged suprahamatus process). Other scapular features of semiaquatic carnivorans indicate strong humeral retraction and abduction thanks to enlarged mm. teres major and acromiodeltoid (e.g. marked caudal angle, concave caudal border, long hamatus process). Unfortunately, a more detailed explanation of the role of these muscles in the locomotion of semiaquatic carnivorans was not possible due to the varying degree of forelimb usage in the studied species (e.g. minks use their forelimbs for both aquatic and overland locomotion, while otters can be considered pelvic oscillators (see above); Williams, 1983; Williams et al., 2002).

#### Climbing

Several shape change trends could be observed in the carnivoran scapula as arboreality increased. Overall, the scapular blade became shorter and wider (Figs. 4, S3). The increase in width was mainly effected by an expansion of the infraspinous fossa, since the width of the supraspinous fossa remained more or less constant except for arboreal species, in which the supraspinous fossa was also enlarged (due to an expansion of the cranial border) (Figs. 4, S3). An expansion of the infraspinous fossa in arboreal species was also described in didelphids (Argot, 2001; Astúa, 2009), while arboreal xenarthrans were characterized by an enlarged supraspinous fossa (Monteiro & Abe, 1999). Additionally, arboreal primates also present large scapular fossae, which have been related to greater shoulder stabilization, relatively heavy limbs, powerful limb movements, and forelimbs used in an extreme protracted or retracted position (Roberts, 1974). As stated above, enlarged fossae are an indication of larger mm. supraspinatus and infraspinatus. The main function of these muscles is shoulder stabilization, but they also provide enhanced mobility of the glenohumeral joint (Roberts, 1974; Argot, 2001), which helps reaching for support in the three-dimensional canopy (Monteiro & Abe, 1999). Another trend observed with increasing arboreality was an increase in the medio-lateral curvature of the scapular blade, as both the vertebral border and, particularly, the neck and glenoid region shifted medially. We hypothesize that this might be related with terrestrial carnivorans often being cursorial (i.e., restricting limb oscillation to the parasagittal plane to optimize running efficiency; Stein & Casinos, 1997), which would benefit from having a flatter scapular blade aligning muscle forces with the rest of the limb. Another potential explanation, maybe complementary, might be that a curved scapular blade would enhance forelimb mobility by allowing a smoother gliding of the scapula relative to the torso.

Regarding the vertebral border, its cranial portion remained more or less constant, while its caudal portion expanded with increasing arboreality (Figs. 4, S3). Thus, the serratus anterior *s. s*. would be larger in arboreal species, providing stronger forelimb retraction, but forelimb protraction would be similar regardless of the degree of arboreality, since the insertion area of the m. levator scapulae remained fairly constant. While the former was expected, since larger forelimb retractors produce larger tensile forces for protracting the body while climbing (Argot, 2001; Astúa, 2009), the latter seemed quite counterintuitive because reaching for a support during arboreal locomotion normally involves extreme forelimb protraction (Roberts, 1974; Monteiro & Abe, 1999). Arboreal didelphids, for instance, also presented enlarged vertebral borders (Argot, 2001), but this was mainly caused by an expansion of its cranial portion instead (i.e., increased forelimb protraction), which was short and angular in terrestrial species. However, the action of enlarged m. supraspinatus of arboreal carnivorans must also be taken into account, since it also acts as a humeral protractor and thus supports the movements of the scapula when reaching for a support. In agreement with this, arboreal didelphids presented relatively smaller supraspinous fossae than terrestrial species (Argot, 2001), since the extra forelimb protraction was not necessary. It is worth noting, however, that Taylor (1974) found differences related to arboreality in the insertion and angulation of the m. levator scapulae of “viverrid-like” carnivorans. Thus, although the general trend in arboreal carnivorans is not to modify this muscle, it has been nonetheless changed in some phyletic lines within Carnivora.

The shape changes of the vertebral border were associated with a cranial displacement of the dorsal end of the scapular spine, especially in arboreal species, in which resulted in a marked curvature of the spine along the dorso-ventral axis (Figs. 4, S3). This increased curvature results in a longer moment arm for the spinodeltoid muscle, which allows a stronger flexion of the shoulder. Similarly, the expansion of the caudal portion of the vertebral border and of the infraspinous fossa also resulted in an increased moment arm of m. teres major (Maynard Smith & Savage, 1956; Monteiro & Abe, 1999), one of the major humeral adductors. Both powerful flexion and adduction of the humerus are important in arboreal mammals, since they help pulling the trunk while climbing (Taylor, 1974; Argot, 2001; Astúa, 2009).

The acromion processes became coplanar with increasing arboreality. Furthermore, as observed in arboreal didelphids (Argot, 2001; Astúa, 2009), the hamatus process both lengthened and shifted cranially, ending up surpassing the glenoid cavity (Figs. 4, S3). A longer hamatus process increases the moment arm of m. acromiodeltoid, which results in a more powerful yet controlled abduction of the shoulder (Taylor, 1974; Argot, 2001). Additionally, a cranially oriented hamatus process both changes the line of action of the acromiodeltoid (so that it contributes to forelimb extension instead of shoulder flexion) and permits abduction of the humerus without colliding with the acromion (Roberts, 1974). It is worth noting that the hamatus process was relatively shorter in arboreal species, since the whole scapula was shortened, but it still extended beyond the glenoid. The shape and orientation of the suprahamatus process remained more or less constant, suggesting that the function of mm. acromiotrapezius and omotransversarius is not affected by arboreality in carnivorans (cf. other forelimb protractors: mm. levator scapulae and supraspinatus).

Finally, the glenoid cavity shifted medially and became more cranially adducted in arboreal carnivorans (Figs. 4, S3). This shift of the glenohumeral joint sets the humerus in a more adducted position, placing the lower arm under the body, which has been shown to increase stability when moving on narrow supports (Schmitt, 2003; Gálvez-López et al., 2011).

#### Digging

The mean scapula shape of semifossorial carnivorans was not statistically significant from that of semiarboreal, scansorial or terrestrial species (Table 6, Fig. 4). This could probably be related to the small number of semifossorial species in our sample, since semifossorial carnivorans can be significantly distinguished from other locomotor types using other forelimb elements (Van Valkenburgh, 1987; Bertram & Biewener, 1990; Gálvez-López, 2021). Nevertheless, some features were characteristic of the scapulae of semifossorial carnivorans. Relative to the terrestrial mean shape, semifossorial species presented a larger infraspinous fossa, a slightly narrower supraspinous fossa (due to the contraction of the cranial border), a lower spine, a more cranially oriented hamatus process, a larger suprahamatus process, and a medially oriented glenoid cavity (Figs. 4, S3). In their study on locomotor adaptations in mammals, Maynard Smith & Savage (1956) stated that fossorial species presented a wide and short scapular blade, a high scapular spine, and a long acromion overhanging the glenoid cavity. None of these adaptations were found in semifossorial carnivorans in the present study. A latter study on fossoriality in rodents (Lehmann, 1963) also reported high scapular spines and, most significantly, the presence of a teres major process at the caudal angle. Similarly, Monteiro & Abe (1999) reported that the shape of the scapula in fossorial xenarthrans (armadillos) was characterized by an expansion of the infraspinous fossa at the origin of the teres major muscle (i.e., at the caudal angle) and also by a relatively longer caudal border. These adaptations relate to an enlarged m. teres major (wide origin) with a long moment arm (caudal border) and, thus, to the powerful limb retraction required for digging with the forelimbs (Hildebrand, 1985). As stated above, semifossorial carnivorans presented similar adaptations of the infraspinous fossa.

### The evolution of scapular morphology in Carnivora

The reconstruction of the ancestral scapula size and shape suggested that extant carnivorans evolved from a medium-sized scansorial ancestor with a scapula similar in shape to that of some genets and mongooses (Figs. 5, 6). In a previous study, Gálvez-López (2021) analyzed the residual means of several morphological variables measured on the appendicular skeleton of a large sample of carnivorans, and arrived to a similar conclusion. The author proposed that the carnivoran ancestor was a forest-dwelling animal, with a similar limb morphology to extant “viverrid-like” taxa (i.e., viverrids, herpestids, etc.) and mixing terrestrial and arboreal adaptations (Gálvez-López, 2021). Both studies are congruent with the placement of “miacids” at the branch leading to extant Carnivora, since recent studies have described a mixed set of adaptations to arboreality and high-speed running in these fossil species (Wesley-Hunt & Flynn, 2005; Spaulding & Flynn, 2009). Other previous studies have proposed that ambulatory species represent the ancestral and unspecialized morphotype in Carnivora (Taylor, 1989; Schutz & Guralnick, 2007). However, comparison with these studies is beyond the scope of the present work, since a proper distinction between generalized and cursorial (*sensu* Stein & Casinos, 1997) carnivorans would require measuring lateral displacements of the limbs during locomotion (e.g. Jenkins, 1971).

#### Caniformia

Regarding the evolution of the carnivoran scapula, the shape of the ancestral caniform (node 2; Fig. 6) was almost identical to that of the carnivoran ancestor (node 1; Fig. 6A), although scapula size was slightly larger (Fig. 5). During caniform evolution, the scapula became larger and wider (nodes 3, 4; Fig. 6), acquiring the basal size and shape from which the ancestors of the large-bodied ursids and pinnipeds diverged (nodes 45 and 39, respectively; Fig. 6E, H). The large scapula sizes reconstructed for basal caniforms agree with the large body mass values estimated by Flynn et al. (2005) for those nodes. However, Finarelli & Flynn (2006) showed that the large ancestral body size of Caniformia were an artifact caused by sampling extant species exclusively. When these authors reconstructed ancestral body sizes including fossil carnivorans in the sample, a small-sized ancestor to Caniformia was the most parsimonious result (Finarelli & Flynn, 2006). Thus, the reconstructed values for ancestral scapula size obtained in the present study must be considered cautiously, since only extant species were measured (due to the scarcity of complete scapulae in the fossil record).

The scapular blade narrowed and lengthened considerably along the branch leading to the ancestor of extant canids (node 49; Fig. 6B), while the cranial edge of the supraspinous fossa bent laterally towards the cranial border and the acromion processes were significantly reduced. All these changes suggest that the canid ancestor was already adapted for running efficiently, as previously suggested by Wang (1993). Similar shape changes occurred during the evolution of the Canini tribe (node 50 onwards), producing even narrower and relative longer scapulae (e.g. *Canis aureus*, Cau; Fig. 6C). A notable deviation from this trend was the re-acquisition of separate hamatus and suprahamatus processes in *Lycalopex* (node 53). On the other hand, shape changes in the scapulae of the Vulpini tribe (node 55 onwards) were small and mostly occurred along the terminal branches. A characteristic feature of vulpine species was a caudal angle projected caudad (e.g. *Vulpes vulpes*, Vvu; Fig. 6D), which increased the moment arm of m. teres major. As explained above, some authors have related this scapular feature to fossoriality (Lehmann, 1963; Monteiro & Abe, 1999). The fact that foxes often bury their captures for latter consumption supports this finding (Wilson & Mittermeier, 2009).

According to its reconstructed scapula size, the ursid ancestor had a similar size than extant pandas (*Ailuropoda melanoleuca*; Fig. 5). However, this must be considered cautiously, since the fossil record suggests that early ursids were small- to medium-sized animals (McLellan & Reiner, 1994; Finarelli & Flynn, 2009). The expanded caudal border of the scapula shape reconstructed for the ursid ancestor (node 45; Fig. 6E) suggests that the enlarged postscapular fossa was already developed. Similarly, the scapula of the ursid ancestor presented an almost rectangular scapular blade and a broad neck. All these scapular features have been previously described by Davis (1949) as characters distinguishing bears from other carnivorans. The scapula shape of the ursid ancestor was characterized by a relatively short and wide scapular blade, an expanded vertebral border (particularly in its caudal portion), and a wide and cranially-oriented acromion extending beyond the glenoid cavity (Fig. 6E). All these scapular features are typical of arboreal carnivorans (see *<u>Climbing</u>*, above), which suggests that the ancestor of extant ursids had a higher degree of arboreality than extant species, as already suggested by Oxnard (1968). All these arboreal features were further developed in the ailurine line, resulting in the extreme scapula shape of the giant panda (*Ailuropoda melanoleuca*, Ame; Fig. 6F). The spectacled bear (*Tremarctos ornatus*, Tor), the only living representative of the tremarctine subfamily, is one of the most arboreal of extant ursids and presents a scapula shape almost identical to the ursid ancestor, although with a longer and more ventrally-oriented acromion (shape not shown). The high degree of arboreality of basal ursids and modern tremarctine bears (Tor) suggests that the alleged adaptations for endurance running of the larger tremarctine species (e.g. *Arctodus*; Matheus, 1997) probably represent an evolutionary dead-end in this lineage. Finally, all ursine bears shared a similar scapula size and shape (node 47 onwards; Figs. 5, 6G), mainly characterized by an extremely convex vertebral border with a marked angulation between its cranial and caudal portions. This scapular feature indicates a clear functional division of the two parts of m. serratus ventralis in protraction and retraction of the forelimb (Maynard Smith & Savage, 1956), and is probably related to the large force required in the particular climbing style of these bears (i.e., “bracing”; Davis, 1949). When “bracing” climbing, the bear first grips the trunk with both forelimbs and pulls the body through the forelimbs (using the forelimb retractors, i.e., m. serratus ventralis *s. s*.) and then secures the hind limbs upper on the trunk while most of the body weight is suspended by the forelimbs (using the forelimb protractors, i.e., m. levator scapulae).

The reconstructed scapula shape for the pinniped ancestor was similar to the mean scapula shape of aquatic carnivorans (node 39; Figs. 4, S3, 6H), supporting the hypothesis that the transition to an aquatic lifestyle occurred shortly after this phyletic line diverged from their shared ancestor with musteloid carnivorans (Arnason et al., 2006; Rybczynski et al., 2009). The main difference between the scapula shape of the pinniped ancestor and the mean shape of aquatic carnivorans was a dorsocaudal expansion of the vertebral border, which is also the scapular feature common to all the oldest pinniped fossils (i.e., *Puijila, Potamotherium, Enaliarctos*; Rybczynski et al., 2009). It is also interesting to note that, in the phylomorphospace defined by PC1 and PC2, the branch leading to Pinnipedia extended almost perpendicularly to the latter at null PC2 values (Fig. 3), indicating that both fossae were enlarged simultaneously prior to the Phocidae/Otariidae split. Then, after the split, otariids expanded towards positive PC2 values (i.e., enlarged supraspinous fossa) and phocids towards negative PC2 values (i.e., enlarged infraspinous fossa. Otariids swim using their forelimbs for propulsion (pectoral oscillators), while in phocids forward thrust is generated with axial undulation and hind limb paddling (pelvic oscillators; see *<u>Swimming</u>*, above). Thus, the expansion of the supraspinous and infraspinous fossae could be related to swimming using the forelimbs or hind limbs, respectively. This finding seems congruent with the hypothesis that early pinnipeds swam quadrupedally using both the forelimbs and the hindlimbs for propulsion (Rybczynski et al., 2009), since both fossae were enlarged simultaneously during the early evolution of this clade. The reconstructed ancestral shapes for the internal nodes of Otariidae and Phocidae further support this interpretation of the phylomorphospace: an expansion of the supraspinous fossa was characteristic of the otariid ancestor (node 40; Fig. 6I), while the infraspinous fossa was enlarged in basal phocids (nodes 43, 44; Fig. 6J). Additionally, a secondary expansion of the supraspinous fossa was observed in the elephant seal (*Mirounga leonina*, Mle; Fig. 6K), probably reflecting the need of powerful forelimb protractors to drag their heavy body on land. Finally, it is worth noting that no significant shape changes occurred during otariid evolution, as it would be expected given the recent radiation of this clade (Nyakatura & Bininda-Emonds, 2012; Fig. 1).

According to its reconstructed scapula size and shape, the musteloid ancestor was a medium-sized scansorial mammal (node 5; Figs. 5, 6L). Its scapula shape was very similar to the carnivoran ancestor (node 1; Fig. 6A), but with a slightly expanded cranial border and a dorsally extended acromion. It also matched closely the mean shape of scansorial carnivorans (Figs. 4, S3), although the musteloid ancestor had a wider scapular blade and neck. Unfortunately, the fossil record for the Musteloidea is incomplete for most lineages, especially regarding the postcranium (Kurtén & Anderson, 1980; Wolsan, 1993). Thus, a direct comparison between the locomotor type inferred from the reconstructed scapula shape and that of early musteloid fossils could not be made. Fabre et al. (2013*b*) conducted a similar study on extant musteloids focusing on the shape of the forelimb long bones, but they reported equivocal results when trying to infer the locomotor type of the musteloid ancestor. The reconstructed centroid size of the musteloid ancestor was 182.93 mm, which corresponded to extant carnivorans in the 4.5 – 9.5 kg body mass range. These values are intermediate between the body mass estimate produced by Finarelli & Flynn (2006) using only extant species and that recovered in the same study after including fossils in the sample. No significant shape changes occurred in the internal nodes representing the divergence of the musteloid families (nodes 5-7; Fig. 6), while reconstructed centroid size values decreased, but remained within the same body mass range (Fig. 5).

The mephitid ancestor presented a particular scapula shape that could not be matched to any of the locomotor type mean shapes (node 38; Figs. 4, S3, 6M). However, it did present several scapular features associated with fossoriality, namely an enlarged infraspinous fossa, a low scapular spine, and a large acromion (particularly the suprahamatus process) (see *<u>Digging</u>*, above). Up to this date, no postcranial elements of early mephitids have been found, and thus nothing is known of their locomotor habits. Regarding scapula size, the reconstructed value suggested a marten-like size for this ancestor (Fig. 5), which is somewhat higher to previous body mass estimations for early mephitids (Finarelli & Flynn, 2009). Finally, it must be noted that only two mephitid species were measured in the present study (Table 1), and thus the reconstructed ancestral scapula size and shape for this clade should be considered cautiously.

Previous studies have suggested an arboreal origin for Procyonidae (Romer, 1966; Baskin, 1982; Fabre et al., 2013*b*), which is strongly supported in the present study. The scapula shape reconstructed for the procyonid ancestor was similar to the mean shape of arboreal carnivorans, but presented even longer and wider acromion processes and a larger scapular blade (node 32; Figs. 4, S3, 6N). The scapular features related to arboreality (e.g. wide and short scapular blade, long and cranially-oriented acromion) were further developed in the most arboreal extant species: the kinkajou (*Potos flavus*, Pfl) and the olingo (*Bassaricyon gabbii*, Bga) (Fig. 6O). Since the kinkajou line diverged early from the procyonid ancestor and the olingos are currently considered sister-taxa to the coatis (*Nasua sp*.; Koepfli et al., 2007), these shape changes must have occurred twice, being more pronounced in the former line. Additionally, in the olingo/coati line the acromion processes were widened (particularly the suprahamatus process) (node 34 onwards; Fig. 6). On the other hand, during the evolution of the raccoon line the scapula became slightly longer and narrower, probably reflecting the less arboreal habits of this clade (node 36 onwards; Fig. 6P).

The reconstructed scapula size and shape for the mustelid ancestor (node 8; Fig. 6) were very similar to that of the musteloid ancestor (node 5; Figs. 5, 6L), indicating that it would also have been a scansorial animal. These results agree with the arboreal habits attributed to *Plesictis*, one of the oldest fossil mustelids, based on its cranial morphology (Palmer, 1999), and also agree with a previous locomotor reconstruction based on extant taxa (Fabre et al., 2013*b*). Both badgers (i.e., *Meles meles*, Mme, and *Mellivora capensis*, Mca) presented highly modified scapula shapes, probably due to their early divergence from the mustelid stem (Fig. 6Q). Both species were considered semifossorial, and as such presented several adaptations to digging (e.g. wide infraspinous fossa, medially oriented glenoid cavity). The rest of the mustelid subfamilies arose in a diversification burst between the middle and late Miocene (Koepfli et al., 2008), which in terms of scapula shape corresponded principally to a reduction of the infraspinous fossa and a shift of the acromion to a more vertically oriented position (nodes 10–13; Fig. 6). Both of these shape changes could be related to a decrease in arboreality (see *<u>Climbing</u>*, above), which would suggest that the arboreal habits were reacquired during the evolution of Martinae (node 26 onwards; Fig. 6R). Indeed, in the internal nodes leading to the extant martens, the hamatus process became progressively longer and more cranially oriented, and the infraspinous fossa expanded again. Another subfamily appearing in this diversification burst was Lutrinae (node 14 onwards; Fig. 6S). However, the shape changes associated to their evolution will not be discussed, since the scapulae in this clade closely matched the mean shape described for semiaquatic carnivorans (Figs. 4, S3), probably because most species in this category were lutrines (Table 1). The shape changes described for the diversification burst were particularly evident in galictine (node 24 onwards; Fig. 6) and musteline species (node 19 onwards; Fig. 6T), in which also an expansion of the cranial border was observed. Overall, the scapula shape of these terrestrial mustelids was quite different to that of other terrestrial carnivorans, and presented features associated with both over ground locomotion (e.g. reduced infraspinous fossa, ventrally directed acromion) and arboreality (e.g. expanded supraspinous fossa, enlarged acromion processes). Schutz & Guralnick (2007) found similar difficulties when trying to infer the locomotor type of *Trigonictis*, a basal mustelid probably related to galictine species, using long bone shape. It could be argued that their characteristic body plan (short legs and long body) would constraint their locomotor performance and that this particular appendicular morphology would be a consequence of it. However, this seems an unlikely explanation, since Horner & Biknevicius (2010) shown that locomotion in terrestrial mustelids is similar to that of other small mammals. On the other hand, their small size could explain these mixed adaptations. As suggested by Astúa (2009), all small mammals could be considered functionally “scansorial” because, for those species, overland and arboreal locomotion pose similar challenges, as some climbing is usually necessary to surpass most obstacles.

#### Feliformia

The earliest feliform fossils belong to the late Eocene and early Oligocene, and already can be ascribed to either the felid or the “viverrid-like” lines (Rose, 2006). Thus, nothing is known of the early evolution of Feliformia, during which the *Nandinia* line and the felid lineage split from the feliform stem (Johnson et al., 2006; Nyakatura & Bininda-Emonds, 2012; Fig. 1). The reconstructed scapula shapes for basal feliforms (nodes 57, 58; Fig. 6) were practically identical to that of the carnivoran ancestor (node 1; Fig. 6A), but with a slightly larger and more cranially directed acromion. According to this, basal feliforms probably would also be scansorial animals, although slightly more arboreal than the earliest carnivorans. On the other hand, scapula size decreased to *c*. 223 mm, which corresponds to the centroid sizes of several extant feliforms in the 10 – 12 kg body mass range (e.g. ocelot, *Leopardus pardalis*; Fig. 5). Thus, it would seem that a size reduction associated with increasing arboreal habits was the main evolutionary trend during early feliform evolution. This trend was probably maintained in the nandiniid line, judging for the scapula size and shape of its small and arboreal lone living representative, the African palm civet (*Nandinia binotata*, Nbi; Fig. 6a).

In agreement with the trend of increasing arboreality proposed for basal Feliformia, the oldest felid, *Proailurus*, was an ocelot-sized carnivoran of the late Oligocene, probably arboreal, since its appendicular skeleton suggests that it was a better climber than most extant species (Agustí & Antón, 2002). Subsequently, felid evolution shifted towards increasingly larger and less arboreal species, a transition that can be observed in the different species of *Pseudaelurus*, a probable descendant of *Proailurus*. The oldest members of *Pseudaelurus* were cat-sized and arboreal, while younger species attained the size of a puma (*Puma concolor*) and were considered more terrestrial (Agustí & Antón, 2002). The reconstructed scapula shape of the ancestor of extant felids suggested a similar locomotor habit, since it was similar to the mean shape of scansorial carnivorans but with a dorsally expanded vertebral border (node 59; Figs. 4, S3, 6b). Additionally, although none of the sampled species presented a similar scapula size than the felid ancestor, the closest values belonged to carnivorans in the puma size range (e.g. *Hyaena hyaena*, Hhy; Fig. 5), and are thus similar-sized to the youngest species of *Pseudaelurus*. According to recent molecular phylogenetic analyses (Johnson et al., 2006), all extant felid species share a common ancestor dating back the late Miocene, which would explain the low shape variability in the scapulae of extant felids (Fig. 6b–e). Nevertheless, significant shape changes could be observed in the scapula during felid evolution, particularly when comparing the two subfamilies. The evolution of pantherine cats was characterized by a ventral displacement of the caudal angle and a dorsal expansion of the vertebral border at its insertion with the scapular spine, which resulted in an increased angulation of the caudal and cranial portions of the vertebral border (node 72 onwards; Fig. 6c). This scapular feature has been related to the functional division of the two parts of m. serratus ventralis in protraction and retraction of the forelimb (Maynard Smith & Savage, 1956). Further shape changes observed in Pantherinae were a reduction of the acromion processes and a caudal expansion of the neck and glenoid region. Similar scapular features developed during the evolution of Ursidae (Fig. 6E, G), suggesting that these might be allometric shape changes. On the other hand, during the evolution of feline cats the caudal portion of the vertebral border expanded dorsally, the scapular neck narrowed, and the acromion processes became larger (e.g. *Felis silvestris*, Fsi; Fig. 6d). Additionally, some shape changes were observed in the terminal taxa of both subfamilies, for instance, an expansion of the fossae at the cranial and caudal angles, and several reorientations of the acromion processes. Finally, it is worth noting that, as specialized runners, both the cheetah and the canids converged in a similar scapula shape (compare *Acinonyx jubatus*, Aju, Fig. 6e, with canids, Fig. 6C, D). To produce such convergence, several shape changes occurred along the branch leading to the cheetah, including a reduction of the supraspinous fossa, a lengthening of the scapular blade and glenoid region, and a ventral shift of the hamatus process.

The remaining extant “viverrid-like” feliforms share a common ancestor dating back to the early Oligocene (Nyakatura & Bininda-Emonds, 2012; Fig. 1). The reconstructed scapula size and shape of this common ancestor (node 76; Figs. 5, 6) suggest that the early evolution of “viverrid-like” carnivorans followed the feliform trend of increasing arboreality and decreasing size. This trend was further continued in the early evolution of Viverridae, since the reconstructed scapula size and shape of the viverrid ancestor indicated that it was civet-sized and semiarboreal (node 77; Figs. 5, 6f). However, along the line leading to extant herpestids, euplerids and hyaenids, the trend shifted towards decreasing arboreality, as suggested by the longer and narrower scapula shape reconstructed for their common ancestor (node 86; Fig. 6i).

The scapula shape of the viverrid ancestor was similar to the mean shape of semiarboreal carnivorans, presenting a wide infraspinous fossa, a cranially oriented acromion, and the characteristic cranial displacement of the dorsal end of the scapular spine (accompanied by the contraction of the cranial portion of the vertebral border, and the expansion of its caudal portion) (node 77; Figs. 4, S3, 6f). These scapular features were further developed in Genettinae (node 84 onwards) and, especially, in Paradoxurinae (node 78 onwards), which includes the most arboreal viverrid species (Taylor, 1974; Wilson & Mittermeier, 2009). It is interesting to note the high degree of convergence in scapula shape between the arboreal paradoxurines (e.g. *Arctictis binturong*, Abi; Fig. 6g) and the arboreal procyonids (Fig. 6O). The evolution of Viverrinae involved a rather different set of shape changes in the scapula. Civets are mostly terrestrial viverrids and thus their scapula shape changed accordingly (e.g. *Viverra zibetha*; Fig. 6h): the supraspinous fossa became smaller and bended laterally, the scapular spine straightened, and the acromion processes ceased to be coplanar (see *<u>Climbing</u>*, above). The earliest viverrid fossils belong to the early Miocene of Eurasia (e.g. *Semigenetta*; Veron, 2010), and have been described as small- to medium-sized scansorial carnivorans (Agustí & Antón, 2002; Morlo et al., 2010). These fossils probably represent basal members of the Genettinae + Viverrinae clade (Veron, 2010), which would explain their less arboreal habits (particularly if they were more related to Viverrinae).

As stated above, herpestids, euplerids and hyaenids share a common ancestor whose scapula shape suggested a more terrestrial habit that basal “viverrid-like” carnivorans (node 86; Figs.4, S3, 6i). A very similar scapula shape was recovered for the common ancestor of herpestids and euplerids (node 87). Besides being longer and narrower than in previous nodes, the scapulae of these internal nodes presented reduced acromion processes and a more laterally curved supraspinous fossa, which are further indicators of decreased arboreality (see *<u>Climbing</u>*, above). Regarding size, although the reconstructed centroid size decreased between these two internal nodes, it was similar to that of several extant carnivorans in the 4 – 11 kg body mass range in both nodes (Fig. 5). The oldest fossils of the Hyaenidae + (Eupleridae + Herpestidae) clade date from the middle Miocene and can already be ascribed to Hyaenidae (*Protictitherium*; Agustí & Antón, 2002). Thus, the reconstructed values for nodes 86 and 87 cannot be compared to the fossil record. However, the fact that similar values of both body size (3 – 10 kg; Morlo et al., 2010) and locomotor habits (generalized terrestrial, Morlo et al., 2010; semiarboreal, Agustí & Antón, 2002) were inferred for *Protictitherium* supports the validity of the present reconstruction.

The early evolution of Hyaenidae is characterized by an increase in both size and terrestrial habits, as evidenced by the sequence *Protictitherium* – *Plioviverrops* – *Thalassictis* (Agustí & Antón, 2002), the latter of which was fully terrestrial and in the 20 kg body mass range (Finarelli & Flynn, 2009). This is congruent with the reconstructed scapula size and shape for the ancestor of extant hyaenids (node 99; Figs. 5, 6j), which suggested that it was terrestrial and somewhat smaller than a striped hyaena (*Hyaena hyaena*, Hhy). Additionally, as observed in canids and the cheetah (Fig. 6C, D, e), the scapula shape of the hyaenid ancestor presented all the scapular features previously described as adaptations to running efficiently (Fig. 6j): long and narrow scapula, reduced fossae with laterally curved margins, highly reduced acromion processes, and a marked angulation between the cranial and caudal portions of the vertebral margin. All these scapular features were further developed in extant hyaenids.

The reconstructed scapula shape of the ancestor of extant herpestids was similar to the mean shape of terrestrial carnivorans (node 88; Figs. 4, S3, 6k). However, the caudal angle was significantly expanded caudad, which indicates well-developed digging abilities (see *<u>Digging</u>*, above). This semifossorial habit was further developed during the evolution of social mongooses (Mungotinae, e.g. *Suricata suricatta*, Ssu; Fig. 6l), since the main shape changes observed in the scapula in this clade were an elongation of both the moment arm of m. teres major (expanded caudal border, dorsally projected caudal angle) and the acromion processes (especially the suprahamatus process). According to previous phylogenetic studies (Nyakatura & Bininda-Emonds, 2012), solitary mongooses underwent a large adaptive radiation at the base of the clade (Fig. 1). Thus, most of the internal nodes for Herpestinae (nodes 88 – 91) presented an almost identical scapula shape, which in turn was very similar to that of the herpestid ancestor. On the other hand, several evolutionary trends could be observed in the terminal herpestine taxa. Both the slender mongoose (*Galerella sanguinea*, Gsa) and the Asian members of *Herpestes* (*H. edwardsi*, Hed, and *H. javanicus*, Hja) presented similar fossorial adaptations as social mongooses (e.g. increased moment arm of m. teres major). Due to its long limbs and proximally located limb musculature, the white-tailed mongoose (*Ichneumia albicauda*, Ial; Fig. 6m) is considered the “most cursorial” herpestid (e.g. Taylor, 1974). Accordingly, its scapula shape was highly convergent with that of other carnivorans adapted to running efficiently (hyaenids, canids,…; Fig. 6C, D, e, j). Finally, the largest herpestine species presented similar-shaped scapulae (e.g. *Herpestes ichneumon*, Hic; Fig. 6n), suggesting that these might be allometric shape changes. Regarding scapula size, the reconstructed centroid size of the herpestid ancestor was similar to that of several extant “viverrid-like” taxa weighting about 2 kg (Fig. 5). As a final remark, no comparison with the fossil record was possible for Herpestidae, since the fossil remains of early herpestids consist mostly of teeth and skull fragments (Agustí & Antón, 2002).

According to its scapula shape, the common ancestor of extant Malagasy carnivorans was very similar to early members of the clade Hyaenidae + (Herpestidae + Eupleridae) (nodes 86–87; Fig. 6i). However, it probably was slightly smaller, since its reconstructed centroid size is similar to that of extant carnivorans in the 2 – 9 kg body mass range (Fig. 5). This small terrestrial ancestor with some climbing abilities arrived to Madagascar from Africa, probably by rafting, which was followed by an adaptive radiation to occupy different niches on the island (Wilson & Mittermeier, 2009). This adaptive radiation is clearly reflected in the variation of scapula shape within this clade. Arboreality was regained along the branch leading to the fossa (*Cryptoprocta ferox*, Cfe; Fig. 6o), which specializes in lemur predation. Consequently, several scapular features related to increased arboreality were developed (e.g. enlarged scapular fossae, shorter scapula, enlarged acromion processes). However, some of these convergent scapular features were brought about by shape changes different than those occuring in most other arboreal carnivorans. For instance, the infraspinous fossa was enlarged mainly by an expansion of the caudal border, while the vertebral border remained practically unchanged. Similarly, the scapula was shortened particularly at the neck and glenoid region, not the scapular blade. These differences could be related to the fact that most other arboreal carnivorans originated from an evolutionary trend towards increased arboreality (e.g. procyonids, viverrids), while the opposite seems to be the case for the clade Hyaenidae + (Herpestidae + Eupleridae). In another euplerid line, now represented by the Malagasy civet (*Fossa fossana*, Ffo), arboreality was further reduced, as evidenced by its long and narrow scapula, convergent with that of other terrestrial carnivorans (shape not shown). Finally, the evolution of Malagasy mongoose was characterized by similar shape changes to those described for true mongooses (Herpestidae): an elongation of the acromion processes (especially the suprahamatus), an expansion of the caudal border, and a dorsal projection of the caudal angle (e.g. *Salanoia concolor*, Sco; Fig. 6p). As in Herpestidae, the reconstructed values for ancestral euplerid nodes could not be compared to the fossil record, since there are no fossil remains of early Malagasy carnivorans (Veron, 2010).

## Acknowledgements

We would like to thank the curators of the Phylogenetisches Museum (Jena), the Museum für Naturkunde (Berlin), the Museu de Ciències Naturals de la Ciutadella (Barcelona), the Muséum National d’Histoire Naturelle (Paris), the Museo Nacional de Ciencias Naturales (Madrid), the Museo Argentino de Ciencias Naturales (Buenos Aires), and the Museo de La Plata (Argentina), for granting us access to their collections. We would also show our appreciation to the following organisations for partially funding this research: la Caixa; Deutscher Akademischer Austausch Dienst (DAAD); the University of Barcelona (UB); Agència de Gestió d’Ajuts Universitaris i de Recerca (AGAUR); Departament d’Innovació, Universitats i Empresa de la Generalitat de Catalunya; and the European Social Fund (ESF). Finally, this work was completed with the assistance of funds from research grants CGL2005-04402/BOS and CGL2008-00832/BOS from the former Ministerio de Educación y Ciencia (MEC) of Spain.

