## Supplementary material for "Evolution of scapula size and shape in Carnivora: locomotor habits and differential shape scaling": S1

**Figure S1. Regressions of shape variables on centroid size (CS).** 95% Confidence ellipses are shown for each family except Ailuridae and Nandiniidae (monotypic families).

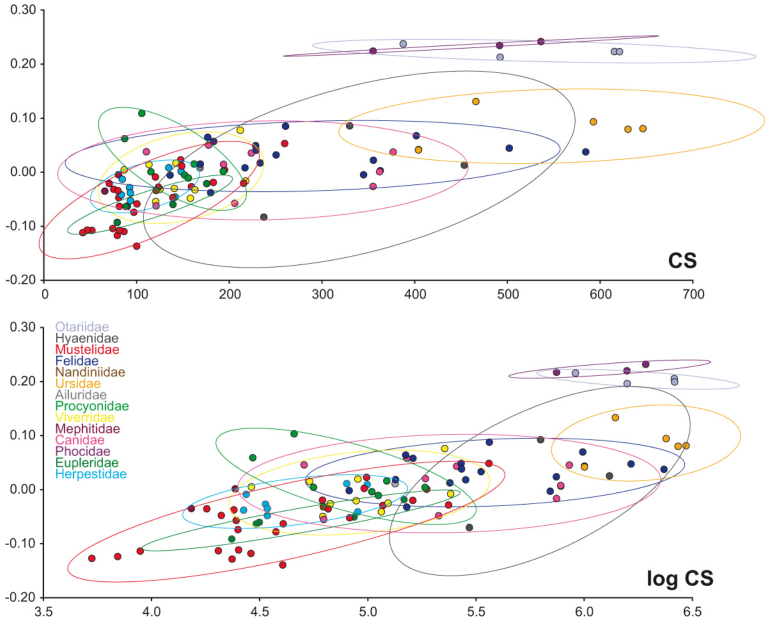


**Figure S2. Principal components analysis of shape variation in the carnivoran scapula, colored by locomotor type.** See Table 2 for definition of the locomotor types, and Figure 3 for the shape changes associated to each principal component.

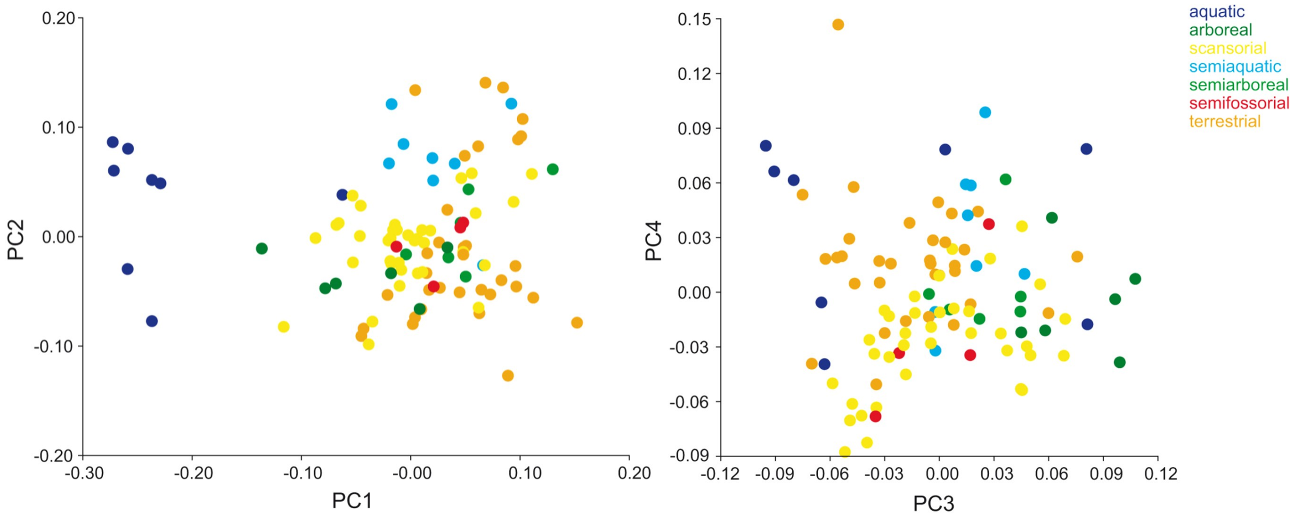

**Table S1. Regressions of shape variables on centroid size in the different subsamples.** The allometric effect on each subsample was determined using regression methods, which allows the shape variation explained by size changes to be expressed as a percentage of the total (between parentheses in the table). Furthermore, the presence of phylogenetic signal was tested on each subsample using permutation tests. See Table 2 for definitions of the locomotor types. Non-significant results (i.e., p-value > 0.05) are presented in ***grey bold italics***. Abbreviations: CS, results of the regression of shape onto centroid size; log CS, results of the regression of shape on log-transformed centroid size; n, sample size; phylogeny, significance of the permutation tests on phylogenetic signal; PIC, results of the regression of shape on size using phylogenetically independent contrasts.

| | **n** | **CS** | **log CS** | **phylogeny** | **PIC** |
|---|---|---|---|---|---|
| **Canidae** | 9 | ***p = 0.8013 (6.78%)*** | ***p = 0.7463 (7.33%)*** | ***p = 0.1396*** | ***p = 0.4993 (8.11%)*** |
| **Ursidae** | 5 | ***p = 0.2128 (33.67%)*** | ***p = 0.2182 (33.11%)*** | p = 0.0143 | ***p = 0.9577 (9.32%)*** |
| **Procyonidae** | 7 | ***p = 0.1076 (31.43%)*** | ***p = 0.0648 (33.75%)*** | ***p = 0.1247*** | ***p = 0.0961 (28.59%)*** |
| **Mustelidae** | 25 | p < 0.0001 (20.58%) | p = 0.0001 (20.93%) | p < 0.0001 | p = 0.0198 (11.41%) |
| **Felidae** | 18 | p = 0.0009 (19.93%) | p = 0.0019 (18.83%) | ***p = 0.0806*** | ***p = 0.1519 (9.04%)*** |
| **Viverridae** | 10 | ***p = 0.3793 (10.50%)*** | ***p = 0.3658 (10.80%)*** | p = 0.0006 | ***p = 0.4404 (9.94%)*** |
| **Eupleridae** | 5 | ***p = 0.0941 (40.39%)*** | ***p = 0.0727 (40.96%)*** | ***p = 0.0648*** | ***p = 0.6280 (17.26%)*** |
| **Herpestidae** | 8 | p = 0.0373 (27.62%) | p = 0.0350 (26.99%) | ***p = 0.0890*** | p = 0.0454 (23.67%) |
| **arboreal** | 6 | ***p = 0.9064 (8.78%)*** | ***p = 0.8517 (9.73%)*** | ***p = 0.1896*** | ***p = 0.3414 (24.81%)*** |
| **semiarboreal** | 6 | ***p = 0.6771 (14.83%)*** | ***p = 0.6784 (14.36%)*** | p = 0.0046 | ***p = 0.4283 (14.45%)*** |
| **scansorial** | 36 | p < 0.0001 (19.37%) | p < 0.0001 (21.68%) | p < 0.0001 | p = 0.0084 (8.23%) |
| **terrestrial** | 33 | p = 0.0001 (20.09%) | p < 0.0001 (28.40%) | p < 0.0001 | ***p = 0.1267 (5.79%)*** |
| **semifossorial** | 4 | p = 0.0410 (50.30%) | p = 0.0431 (50.03%) | ***p = 0.3311*** | ***p = 0.3318 (51.94%)*** |
| **semiaquatic** | 8 | p = 0.0220 (27.76%) | p = 0.0074 (29.05%) | p = 0.0242 | ***p = 0.2489 (19.43%)*** |
| **aquatic** | 8 | ***p = 0.5356 (12.28%)*** | ***p = 0.4327 (14.34%)*** | p = 0.0105 | ***p = 0.9417 (6.60%)*** |

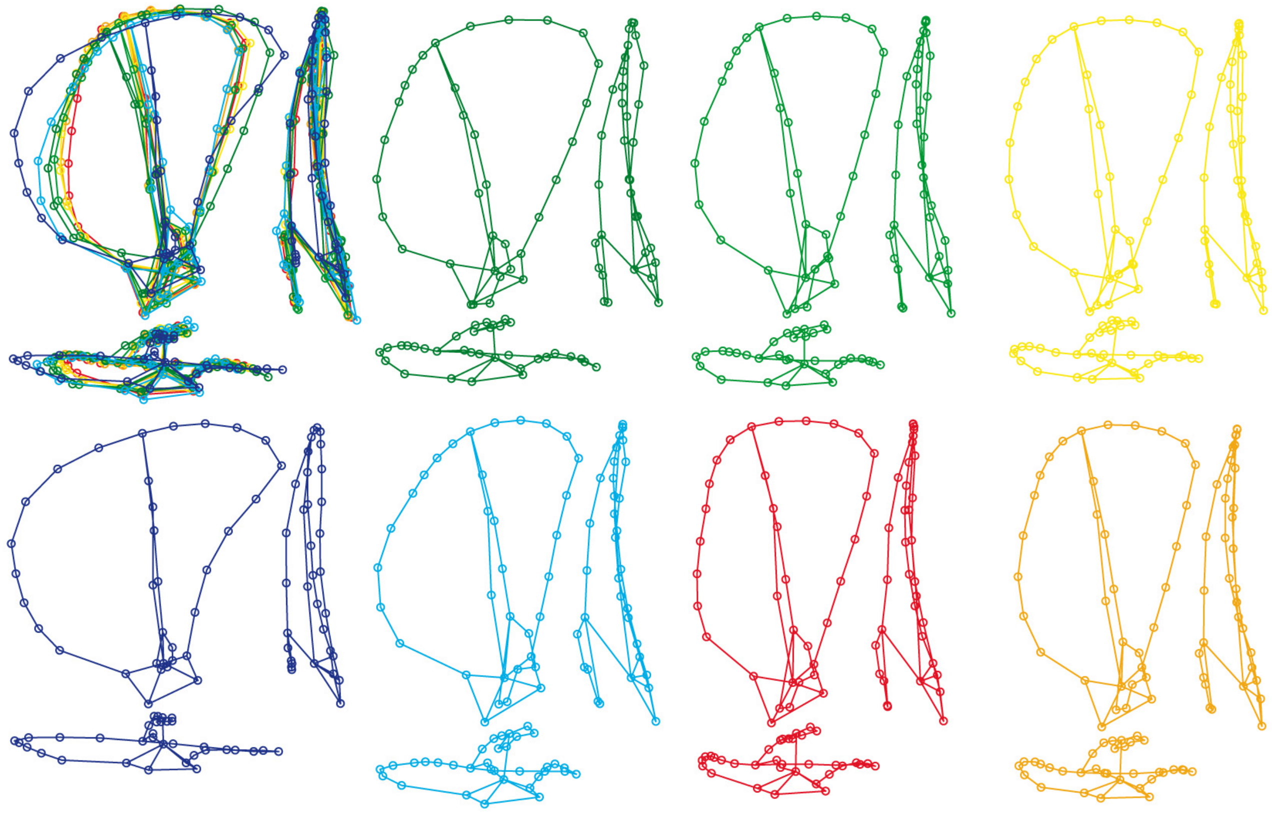

**Figure S3. Mean scapular shape by locomotor type.** For each locomotor type, a set of three wire-frames is presented so that scapular shape can be observed in lateral (x-y; left), dorsal (x-z; bottom), and caudal (y-z; right) views. Furthermore, a superimposition of all mean shapes is presented to ease comparison. See Table 2 for definition of the locomotor types. Locomotor type colors follow the legend in Figure S2: dark green for arboreal, green for semiarboreal, yellow for scansorial, dark blue for aquatic, blue for aquatic, red for semifossorial, and orange for terrestrial.
